# Magnesium depletion unleashes two unusual modes of colistin resistance with different fitness costs

**DOI:** 10.1101/2024.10.15.618514

**Authors:** Yu-Ying Phoebe Hsieh, Ian P. O’Keefe, Zeqi Wang, Wanting Sun, Hyojik Yang, Linda M. Vu, Robert K. Ernst, Ajai A. Dandekar, Harmit S. Malik

## Abstract

Increasing bacterial resistance to colistin, a vital last-resort antibiotic, is an urgent challenge. We previously reported that magnesium sequestration by *Candida albicans* enables *Pseudomonas aeruginosa* to become colistin-resistant. Here, we show that Mg²⁺ depletion drives *P. aeruginosa* to evolve greater colistin resistance through genetic changes in lipid A biosynthesis-modification pathways and a putative magnesium transporter. These mutations synergize with the Mg^2+^-sensing PhoPQ two-component signaling system to remodel lipid A structures of the bacterial outer membrane in previously uncharacterized ways. One predominant mutational pathway relies on early mutations in *htrB2*, a non-essential gene involved in lipid A biosynthesis, which enhances resistance but compromises outer membrane integrity, resulting in fitness costs and increased susceptibility to other antibiotics. A second pathway achieves increased colistin resistance independently of *htrB2* mutations without compromising membrane integrity. In both cases, reduced binding of colistin to the bacterial membrane underlies resistance. Our findings reveal that Mg^2+^ scarcity unleashes two novel trajectories of colistin resistance evolution in *P. aeruginosa*. (160)

## Introduction

Antimicrobial resistance poses a significant global health challenge [1]. Prior research has focused on mechanisms that individual bacterial species use to evade antibiotics. However, microbial interactions can profoundly alter antibiotic resistance in ways that remain incompletely understood [2, 3]. Complex microbial communities can protect susceptible species from antibiotics, modulate selection pressures [4, 5], influence the emergence of resistant mutants [6], and alter the spectrum of resistance mutations [7]. Under these conditions, microbes must continuously adapt to complex selective pressures imposed both by antibiotics and their surrounding microbial partners, making it challenging to predict the evolutionary trajectory of antibiotic resistance. Since polymicrobial infections can exacerbate the spread of drug-resistant bacteria and pose a significant threat to human health and healthcare systems [8–10], understanding how microbial ecology shapes the evolution of antibiotic resistance is critical for developing effective therapies against polymicrobial infections.

*Pseudomonas aeruginosa* is a Gram-negative bacterium commonly found in polymicrobial, drug-resistant infections [11, 12]. Polymyxins, such as colistin (polymyxin E), are a last resort for treating multidrug-resistant *P. aeruginosa* infections [13, 14]. In recent years, the number of colistin-resistant *P. aeruginosa* isolates has increased at an alarming rate [15]. Colistin targets Gram-negative bacteria by electrostatic interaction with lipid A, the hydrophobic membrane anchor of lipopolysaccharide (LPS) in the outer leaflet of the outer bacterial membrane [16, 17], eventually leading to membrane rupture and cell lysis [18]. LPS is the predominant constituent of the outer leaflet of the outer membrane that protects bacteria from environmental changes [19–21]. Bacteria can rapidly modify lipid A to facilitate adaptation to environmental perturbations [22, 23]. For instance, upon exposure to colistin or Mg^2+^ depletion, *P. aeruginosa* activates the PhoPQ and PmrAB two-component systems [24, 25], which activate the expression of several lipid A-modifying proteins, including PagL and proteins encoded by the Arn operon. PagL alters the acyl chain number of lipid A by removing the acyl chain from the 3-position of lipid A [26]. Activation of Arn operons leads to the addition of 2-amino-2-hydroxy-L-arabinose (L-Ara4N) modification onto lipid A [27] [28, 29] (Fig. S1A-1B). These lipid A modifications reduce colistin binding to the outer membrane, resulting in protection from its antibacterial action [25, 30, 31]. Such lipid A modifications are conserved in Gram-negative bacteria [31, 32]. Yet, how polymicrobial environments alter conditions and mechanisms by which bacteria acquire colistin resistance remains underexplored.

*P. aeruginosa* coexists with the fungal pathogen *Candida albicans* in urinary tracts, chronic wounds, and the airways of people with cystic fibrosis [33–36]. In a previous study, we showed that *C. albicans* (and many fungi) sequester Mg^2+^ from *P. aeruginosa* (and many Gram-negative bacteria), which alters the evolutionary trajectory of colistin resistance acquisition by *P. aeruginosa* [37]. In this prior study, we grew eight replicate populations of wild-type (WT) *P. aeruginosa* strain PAO1 in co-culture with *C. albicans* in brain heart infusion (BHI) media with gradually increasing colistin concentration. We found that increased resistance to colistin always depended on either fungal co-culture or Mg^2+^ depletion, as fungal removal or Mg^2+^ supplementation reduced colistin resistance in all replicate populations [37]. Intriguingly, replicate populations that developed increased colistin resistance under high Mg^2+^ conditions acquired an almost entirely distinct set of mutations [37].

Our findings suggested that the low Mg^2+^ conditions experienced by bacteria in polymicrobial environments [38] and during infection [39] can drive alternative modes of colistin resistance. However, the molecular nature of these mechanisms remained unclear. Here, we characterized the genetic, evolutionary, and biochemical mechanisms by which these mutations give rise to novel modes of colistin resistance and investigated why these mutations evolve exclusively under low Mg^2+^ conditions. Our analyses reveal that colistin resistance arises via genetic epistasis between early-arising mutations that synergize with PhoPQ signaling, which is activated under low Mg^2+^ conditions. Furthermore, we demonstrate that these mutations result in distinct lipid A modifications, which confer colistin resistance by reducing colistin binding. Interestingly, many of these changes compromise bacterial membrane integrity, leaving colistin-resistant populations more susceptible to the action of other antibiotics. Our study offers new molecular insights into how the nutritional depletion of metal ions can impact bacterial antibiotic resistance.

## Results

### *P. aeruginosa* replicate populations acquire high levels of colistin resistance under low Mg^2+^ conditions via genetic epistasis

We evolved WT *P. aeruginosa* to gain colistin resistance in co-culture with *C. albicans* by gradually increasing colistin concentration from 1.5 to 192 μg/ml during daily transfers (Fig. 1A) [37]. Our evolution experiments revealed three evolutionary trajectories leading to low Mg^2+^-dependent colistin resistance, with convergent mutations in genes involved in lipid A biosynthesis and modification (Fig. S1A) and a gene encoding a novel Mg^2+^ transporter, *PA4824*. The first trajectory (I) acquired mutations in *htrB2*, *PA4824*, *oprH*, *fimX*, and *PA3647*. The second trajectory (II) acquired mutations in *lpxO2*, *htrB2*, and *PA4824*. The third trajectory (III) acquired mutations in *lpxA*, *PA4824*, *oprH*, *PA5005*, *pilB*, *colS*, and *ftsY*.

**Figure 1.**
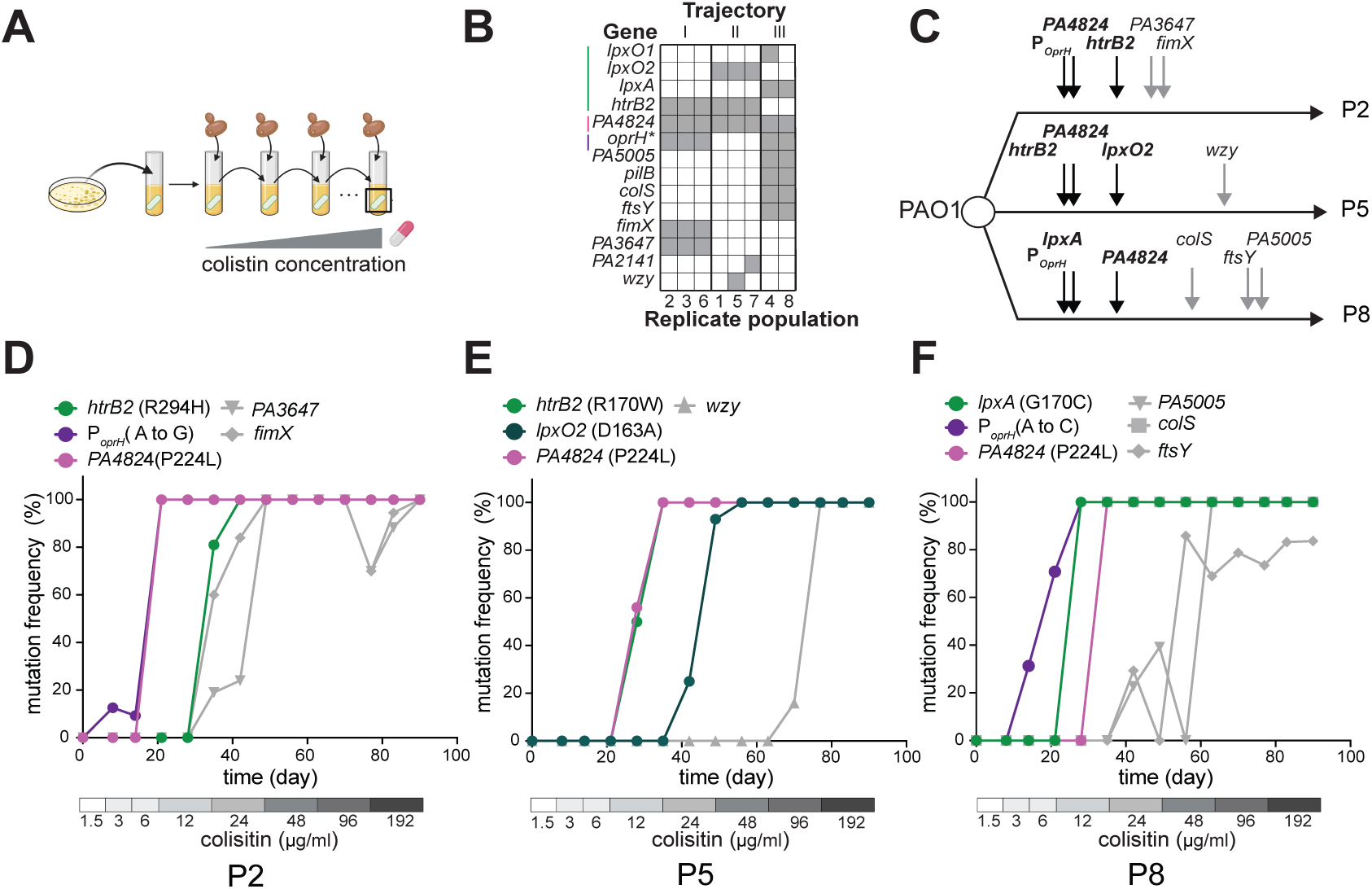
Three distinct evolutionary trajectories lead to fungal-dependent colistin resistance under low Mg^2+^ conditions. **(A)** Schematic of evolving colistin-resistant *P. aeruginosa* in co-culture with *C. albicans*. Eight independent *P. aeruginosa* populations were passaged in BHI media with *C. albicans* (which sequesters Mg^2+^) and colistin. *C. albicans* cells were added at each passage, and the concentration of colistin was gradually increased from 1.5 μg/ml to 192 μg/ml. **(B)** Co-culture evolved populations exhibit three patterns of genetic mutations. Genes mutated in the eight populations are listed in rows, and mutations in a population are indicated as gray boxes (*indicates a non-coding mutation in the promoter of the *oprH*/*phoP*/*phoQ* operon). The eight populations are grouped based on the similarity of their mutation patterns and shown as trajectory I, II, and III. **(C)** Summary of the order of mutation fixation in representative populations P2, P5, and P8. Mutations in lipid A biosynthesis genes, *PA4824*, and the promoter of the *oprH*/*phoP*/*phoQ* operon (labeled in black) were fixed in the early stage of evolution across three trajectories. **(D-F)** Temporal dynamics of mutation fixation in evolved populations P2 (D), P5 (E), and P8 (F) show that mutations in lipid A biosynthesis genes (*htrB2*, *lpxO2*, and *lpxA*), *PA4824*, and the *oprH*/*phoP*/*phoQ* promoter became fixed early during adaptation, when populations were exposed to 12-24 μg/ml colistin.

To determine how these trajectories developed colistin resistance, we used representative isolates from each of the three evolutionary trajectories: replicate populations P2 (from trajectory I), P5 (from trajectory II), and P8 (from trajectory III) (Fig. 1B). Initially, we determined the order and timing of the acquired mutations by performing whole-genome sequencing of samples collected every two weeks during the evolution of each replicate population. We found that missense mutations in lipid A biosynthesis genes (*htrB2*, *lpxO2*, and *lpxA*), the putative Mg^2+^ transporter *PA4824*, and the promoter of the *oprH/phoP/phoQ* operon, arose and were fixed relatively early during exposure to increasing colistin levels (Fig. 1C). For example, in replicate P2, *oprH*/*phoP*/*phoQ* and *PA4824* (P224L) mutations were fixed by day 21 (at 6 μg/ml colistin), while the *htrB2* (R294H) mutation was fixed by day 35 (at 12 μg/ml colistin) (Fig. 1D). In replicate P5, *htrB2* (R170W) and *PA4824* (P224L) mutations came to fixation in the population by day 28 (at 12 μg/ml colistin), while the *lpxO2* (D163A) mutation was fixed by day 42 (at 24 μg/ml colistin) (Fig. 1E). Finally, in replicate P8, the *oprH/phoP/phoQ* promoter mutation arose first and were fixed in the population with the *lpxA* (G170C) mutation by day 28 (at 12 μg/ml colistin), followed by the *PA4824* (P224L) mutation being fixed by day 35 (also at 12 μg/ml colistin) (Fig. 1F). This pattern of convergent evolution – mutations occurring in the same genes across different replicate populations – suggested that these mutations are likely to be causal for acquiring colistin resistance under low Mg^2+^ conditions.

We next investigated how these mutations contribute to colistin resistance under low Mg^2+^ conditions. We used *C. albicans*-spent BHI media, the supernatant from a fungal-saturated culture, as a low Mg^2+^ condition, in contrast to standard BHI media (which has high Mg^2+^) [37]. To assess their causal role in colistin resistance, we reconstituted single, double, and triple mutants – reflecting their temporal emergence in representative P2, P5, and P8 populations – into WT PAO1 to mimic the early stages of bacterial adaptation to increasing colistin concentrations. We then measured the colistin minimal inhibitory concentration (MIC) of these mutants in low Mg^2+^. Combining the mutations we identified as fixed by day 35 in each trajectory was sufficient to confer significant colistin resistance relative to the WT (Fig. 2A-C). In all cases, triple-mutation-reconstructed strains were resistant to colistin concentrations equivalent to or higher than those used at the time points when these mutations were fixed. Consistent with our MIC estimates, triple-early mutation strains had significantly increased survival at colistin concentrations between 6 to 24 μg/ml in co-culture with *C. albicans* (Fig. S2), indicating that the MIC measurements in the low Mg^2+^ media are a reasonable proxy for increased resistance in our co-culture evolution experiments.

**Figure 2.**
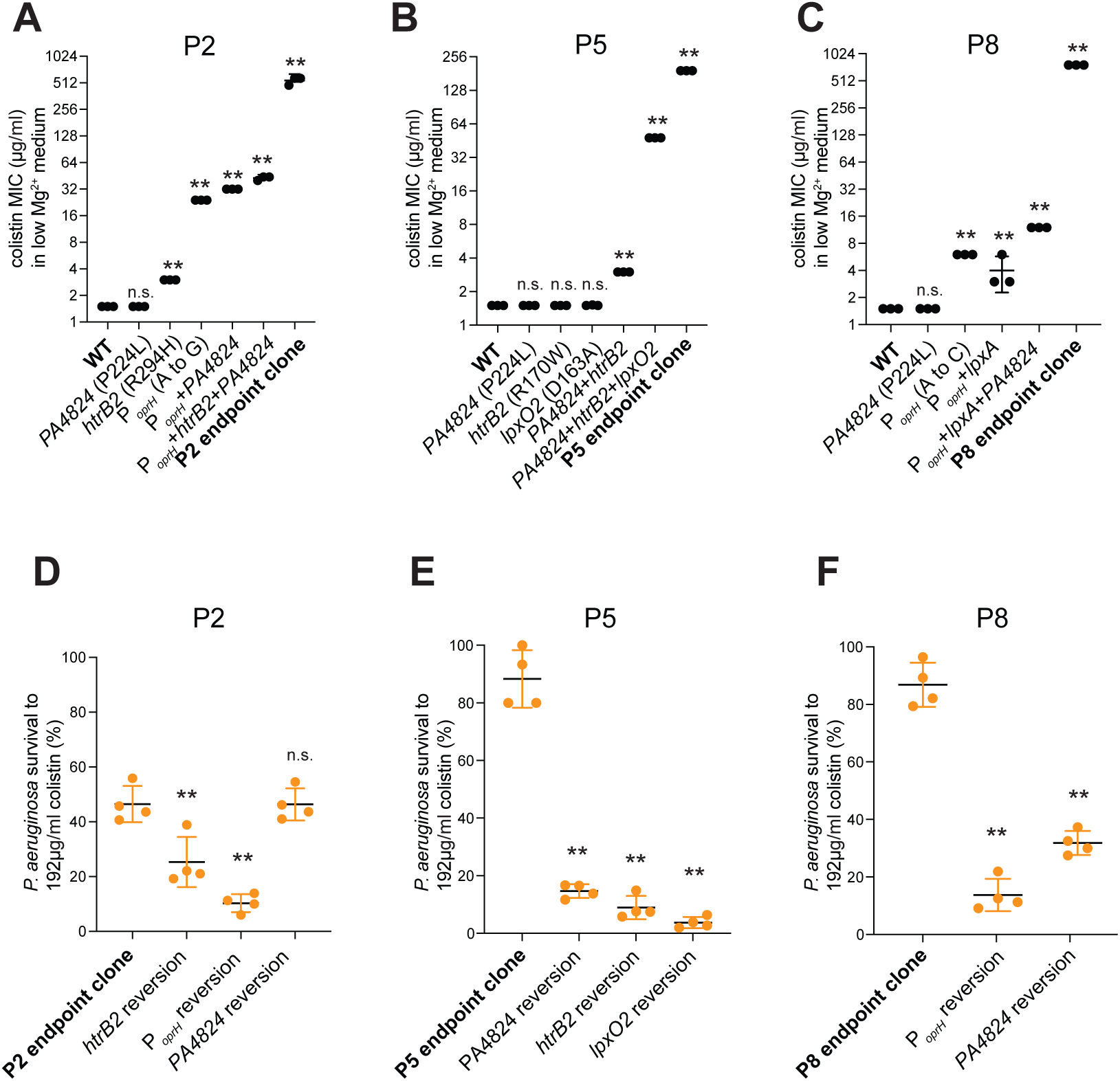
Mutations in lipid A synthetic genes, *PA4824*, and the *oprH*/*phoP*/*phoQ* promoter are necessary and sufficient to explain colistin resistance. **(A-C)** Early-occurring mutations from representative populations P2 (A), P5 (B), and P8 (C) were reconstructed into the WT PAO1 strain. Strains with triple reconstructed mutations, but not single mutations, showed increased colistin MIC compared to WT PAO1 in low Mg^2+^ media, with higher resistance than the levels against which they were selected during the experimental evolution. In all three representative populations, MICs of final evolved endpoint clones were higher than those of the early triple mutant reconstructed strains. Mean ± std of 3 biological replicates is shown (\*\**p* < 0.01, Dunnett’s one-way ANOVA test). **(D)** Reversion of *htrB2* or *oprH*/*phoP*/*phoQ* promoter mutation to the WT allele in a P2 endpoint clone reduced survival in192 μg/ml colistin in the low Mg^2+^ media. In contrast, reversion of PA4824 had no significant effect. **(E)** Reversion of *PA4824*, *htrB2*, or *lpxO2* mutations in a final P5 clone lowered survival to 192 μg/ml colistin in low Mg^2+^ media. **(F)** Reversion of *PA4824* or *oprH*/*phoP*/*phoQ* promoter mutations in a final P8 clone lowered survival in 192 μg/ml colistin in low Mg^2+^ media. Mean ± std of 4 biological replicates is shown. (\*\**p* < 0.01, Dunnett’s one-way ANOVA test)

Strains harboring single or double mutants exhibited a lower degree of colistin resistance compared to the triple-mutant strains, indicating that positive epistasis played a crucial role in the evolution of colistin resistance. This phenomenon is most dramatically observed in P5, where no single mutation (*oprH*, *PA4824, htrB2)* had a discernible effect on colistin resistance, whereas a double mutant (*htrB2+PA4824*) was modestly resistant, and the triple mutant (*htrB2+PA4824*+*lpxO2*) was highly resistant (Fig. 2B). The MIC of the P5-derived triple mutant (48 μg/ml) was fourteen-fold higher than would be expected from the addition of single mutants, highlighting the role of positive epistasis in the P5 population. In contrast to P5, individual *oprH/phoP/phoQ* promoter mutations were sufficient to confer significant colistin resistance to the P2 and P8 populations, which could be further augmented by additional mutations (Fig. 2A, 2C). Intriguingly, although the same *PA4824* mutation (P224L) was found recurrently in all three representative populations, this mutation did not confer any degree of colistin resistance on its own.

While early triple mutations explain the initial emergence of colistin resistance, they were still insufficient to confer resistance at the highest colistin concentrations used in our experimental evolution (192 µg/ml), indicating that subsequent mutations in each trajectory were also essential for maximal resistance (Fig. 2A-C). Nevertheless, we reasoned that early-occurring mutations may have been necessary for the high degree of colistin resistance. To test this hypothesis, we selected a single endpoint clone for each population that encoded all (shared) fixed mutations but encoded the smallest number of unshared mutations in the population. For each of the highly resistant endpoint clones from P2, P5, and P8, we reverted the early-occurring mutations. We then tested their survival when challenged with 192 µg/ml colistin in low Mg^2+^ by measuring colony-forming units. We found that reverting mutations in *htrB2*, *lpxO2*, or the *oprH/phoP/phoQ* promoter significantly reduced bacterial survival. Reverting the *htrB2* mutation to the WT allele in P2 and P5 reduced bacterial survival to ∼20% or less (Fig. 2D-E). Similarly, reverting the *oprH/phoP/phoQ* promoter mutation reduced bacterial survival to 20% or 10% in P2 and P8, respectively (Fig. 2D, 2F), while reverting the *lpxO2* mutation reduced P5 survival to 3% (Fig. 2E). In contrast, reverting the *PA4824* mutation significantly lowered bacterial survival in P5 and P8, but not P2 (Fig. 2D-F), suggesting that other mutations might compensate for *PA4824* evolution in P2. We also found that the colistin MIC was reduced in these reversion strains, following a similar trend as the colistin survival assay (Fig. S3). Our findings establish the cadence and causality of the genetic mutations observed in three distinct evolutionary trajectories. They reveal that “early” mutations in *htrB2*, *lpxO2*, *oprH/phoP/phoQ* promoter, and *PA4824* were sufficient for evolving initial low colistin resistance and are necessary for maintaining high colistin resistance under low Mg^2+^ conditions.

### Novel, distinct lipid A structures underlie low Mg^2+^-dependent colistin resistance

Our experimental evolution uncovered mutations that might directly or indirectly affect lipid A structures (Fig. 1B and S1A). For example, *htrB2* is mutated in two of three evolutionary trajectories, while other lipid A biosynthesis genes (*lpxO1*, *lpxO2*, and *lpxA*) are mutated in at least one trajectory (Fig. 1B). To determine the impact of these mutations on lipid A, we used the fast lipid analysis technique (FLAT) paired with matrix-assisted laser desorption/ionization time-of-flight (MALDI-TOF) mass spectrometry (MS) [40] to characterize the lipid A structures of all replicate populations and compared them to WT PAO1.

WT PAO1 grown in BHI (high Mg^2+^), synthesized expected lipid A structures, with penta-acylated lipid A (m/z=1445.86), which varies in 2-hydroxylation status (m/z=1461.85) and phosphorylation status (m/z=1365.89, 1525.82) (Fig. 3A and S4A) [26, 41]. In contrast, PAO1 grown in fungal-spent BHI (low Mg^2+^) synthesized lipid A that was hexa-acylated, with PagP-mediated palmitate addition (m/z=1684, 1700) [42] or a single L-Ara4N modification (m/z=1576, 1592) [43] (Fig. 3B and S4B), which are mediated by the PhoPQ and PmrAB two-component systems [24, 25]. In contrast to WT PAO1, all eight colistin-resistant populations (P1 through P8) demonstrated mass spectra that are unique to each evolutionary trajectory, and distinct from the WT in both high and low Mg2^+^ conditions (Fig. S5). Intriguingly, these alterations were observed at both high and low Mg^2+^ conditions (Fig. S5), suggesting that they were genetically determined and not induced by low Mg^2+^ conditions.

**Figure 3.**
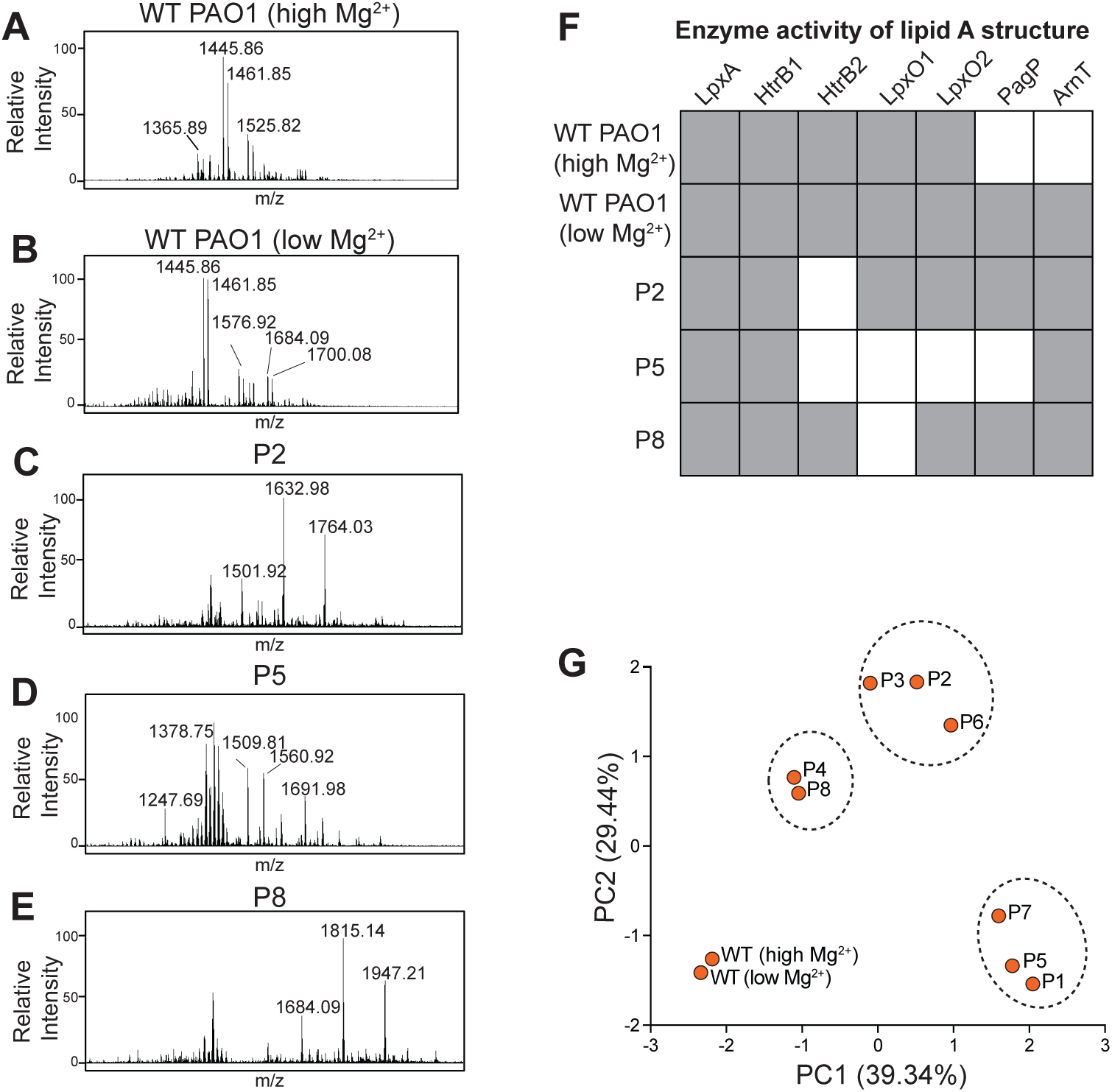
Novel, distinct lipid A structures in each of three evolutionary trajectories of low Mg^2+^-dependent colistin-resistant populations. **(A)** Mass spectra of WT PAO1 in high Mg^2+^ media reveal major ions at m/z values 1365.89, 1445.86, 1461.85, 1525.82, and 1451.82. This lipid A is penta-acylated [26], with both single and double 2’-hydroxylation mediated by LpxO1/LpxO2 [41], indicated by differences of approximately 16 m/z, and single, double, and triple phosphorylation status, indicated by differences in approximately 80 m/z from the base peak. **(B)** Mass spectra of WT PAO1 in low Mg^2+^ media reveal additional hexa-acylated lipid A with PagP-mediated palmitoylation [42], as seen by ions 1684.09 and 1700.08, or single L-Ara4N addition from the base penta-acylated structure [43], shown by the ion at m/z=1576.92. **(C)** Mass spectra analyses of P2 cells reveal penta-acylated lipid A with PagP-mediated palmitate addition but lacking HtrB2-mediated laurate addition, leading to *m/z* values of 1501.01, 1632.98, and 1764.03. **(D)** Mass spectra analyses of P5 cells reveal lipid A either lacking HtrB2-mediated acylation and LpxO2-mediated 2-hydroxylation (*m/z* = 1378.75) or containing HtrB2-mediated acylation but lacking both LpxO1 and LpxO2-mediated 2-hydroxylation (*m/z* = 1560.92). **(E)** Mass spectra analyses of P8 reveal hexa-acylated lipid A with PagP-mediated palmitoylation but lacking LpxO1-mediated hydroxylation (*m/z* = 1684.09, 1815.14, 1947.21). In addition, lipid A in P2, P5, and P8 cells exhibits single or double L-Ara-4-N addition, shown by differences of approximately 131 *m/z*. **(F)** A summary table of the activity of lipid A biosynthesis and modification enzymes. Lipid A structures obtained from MALDI-TOF MS were used to infer the activity of each enzyme (gray box indicates functional enzyme). The WT strain and endpoint clones (P2, P5, and P8) were analyzed in high and low Mg2^+^ conditions. In the endpoint clones, the activity of these enzymes was similar in both conditions. **(G)** PCA of lipid A mass spectra from all eight evolved populations and the laboratory-adapted WT strain PAO1 based on the presence or absence of lipid A peaks. PC1 and PC2 are shown, with 39.34% and 29.44% of the variance explained, respectively. Ellipses group strains that cluster together based on observed lipid A structures from FLAT.

Our analysis identified several lipid A structures in each of the three evolutionary trajectories that have not been previously reported in *P. aeruginosa*, including those in colistin-resistant strains [44, 45]. These distinct lipid A structures include species with various acylation and L-Ara4N modifications. For example, populations P2, P3, and P6 (trajectory I) synthesized penta-acylated lipid A with PagP-mediated palmitate addition but lacked HtrB2-mediated laurate addition (m/z=1632.96) (Fig. 3C and S4C, Table S1). In contrast, populations P1, P5, and P7 (trajectory II) exhibited mixtures of two distinct lipid A structures without PagP-mediated palmitate addition. These trajectory II variants exhibited tetra-acylated lipid A that lacked HtrB2-mediated acylation and lpxO2-mediated hydroxylation (*m/z* = 1378.75), and penta-acylated lipid A with HtrB2-mediated laurate addition, but without LpxO1 and LpxO2-mediated hydroxylation (*m/z* = 1560.92) (Fig. 3D and S4D, Table S1). Finally, populations P4 and P8 (trajectory III) had hexa-acylated lipid A containing PagP-mediated palmitate addition but lacking LpxO1-mediated hydroxylation (*m/z* = 1815.14) (Fig. 3E and S4E, Table S1). The lipid A of these evolved populations had absent, single or double L-Ara4N addition, shown by differences of approximately 131 m/z between ions in Fig. 3C-3E. These results demonstrate that selection for colistin resistance in low Mg^2+^ leads to dramatic diversification of lipid A structures (Fig. 3F and S4). Principal component analysis (PCA) of the MS data revealed that lipid A structures of the eight evolved populations formed three separate clusters (Fig. 3G), each distinct from WT PAO1, mirroring the three evolutionary trajectories we had observed previously (Fig. 1B).

To identify which genetic mutations underlie these biochemical changes, we analyzed early-occurring triple mutants (described above). We found that early-occurring mutations in P2 and P5 can fully recapitulate the lipid A structures of endpoint clones (Fig. S6A-6B), whereas triple mutants of P8 partially recapitulate the final lipid A structures, still containing LpxO1-mediated 2’-hydroxylation (Fig. S6C). Since these early-occurring triple mutants also showed increased resistance compared to the ancestral strain (Fig. 2A-2C), we conclude that the unique lipid A changes associated with these early mutations causally contributed to colistin resistance.

We also assessed the necessity of specific mutations for the distinct lipid A structures found in each trajectory by performing FLAT on each mutant-reversion strain. Reverting individual mutations in *htrB2* or *lpxO2* to the wild type (WT) restored the laurate moiety (Fig. S7A and S7D) or 2’-hydroxylation of lipid A (Fig. S7F), respectively, indicating that these mutations disrupt enzyme function. Reversion of *oprH/phoP/phoQ* promoter mutations prevented the addition of PagP-mediated palmitoylation (Fig. S7C and S7G), a modification activated by PhoPQ. This result suggests that these promoter mutations affect downstream effectors of the PhoPQ pathway. These results indicate that, despite their importance to lipid A biosynthesis in *P. aeruginosa* [46, 47], loss-of-function mutations in *htrB2* and *lpxO2* can nevertheless increase colistin resistance under low Mg^2+^ conditions by altering lipid A structures in novel ways.

In contrast, mutations in *PA4824* and *lpxA* appear to contribute to colistin resistance without altering lipid A. Reversion of the *PA4824* mutation (P224L) in all three endpoint clones had no discernible effect on lipid A structures (Fig. S7B, S7D, and S7F). LpxA is the first, essential enzyme in lipid A biosynthesis (Fig. S1) [48]. Despite the same *lpxA* mutation (G170C) being found in two populations of trajectory III, P8 endpoint clones had LpxA-mediated acylation in their lipid A structures, indicating this mutation does not impair the enzymatic function of LpxA.

Overall, our findings revealed that high colistin resistance in low Mg^2+^ conditions depends on unique lipid A structures that arise via two distinct mechanisms: loss-of-function mutations in *htrB2* and *lpxO2*, and promoter mutations in the *oprH/phoP/phoQ* operon. In contrast, mutations in *PA4824* and *lpxA* drive higher colistin resistance independent of changes in lipid A structures.

### The PhoPQ pathway synergizes with early-arising mutations to potentiate colistin resistance under low Mg^2+^ conditions

We next investigated why the mutations conferring robust colistin resistance in low Mg^2+^ conditions are not observed in Mg^2+^ replete conditions. Consistent with many prior studies [28, 29], we found that bacteria activate the PhoPQ and the PmrAB two-component systems to modify their membrane in low Mg^2+^ (Fig. S8). We hypothesized that such physiological changes might interact synergistically with early-occurring mutations to enhance colistin resistance in low Mg^2+^. Moreover, two of the evolutionary lineages (represented by P2 and P8) had distinct mutations upstream of the *oprH*/*phoP*/*phoQ* operon (Fig. 1B). To distinguish whether these promoter mutations acted via OprH and/or PhoPQ to enhance colistin resistance, we deleted either *oprH* or *phoP* in endpoint clones from P2 and P8. In both cases, the deletion of *phoP,* but not *oprH,* significantly decreased the colistin MIC in the low Mg^2+^ media (Fig. S9). These results suggested that the promoter mutations were selected for their ability to activate PhoPQ and not OprH.

To test whether *oprH*/*phoP*/*phoQ* promoter mutations could alter the expression of PhoPQ-regulated genes, we introduced each mutation into the WT strain and measured the expression of two known PhoPQ targets: *arnC* [49] and *pagP* [50]. In both high and low Mg^2+^ media, we found that *arnC* and *pagP* were upregulated in the promoter-mutation-reconstructed strains relative to WT (Fig. 4A and 4B). Using lipid A structural analysis, we also found that both *phoPQ* promoter mutations introduced into WT PAO1 were sufficient to activate PhoPQ-mediated lipid A modifications that are typically observed in low Mg^2+^ conditions, even in high Mg^2+^, recapitulating the same L-Ara4N modification and PagP-mediated acylation observed in populations P2 and P8 (Fig. 4C-4E). In contrast, loss of *phoP* in P2 and P8 endpoint clones completely abrogated L-Ara4N modification and PagP-mediated acylation of LPS (Fig. S10), consistent with previous reports [24, 30]. Collectively, our data confirms that each promoter mutation is sufficient to drive lipid A modifications that enhance colistin resistance independent of Mg^2+^ levels.

**Figure 4.**
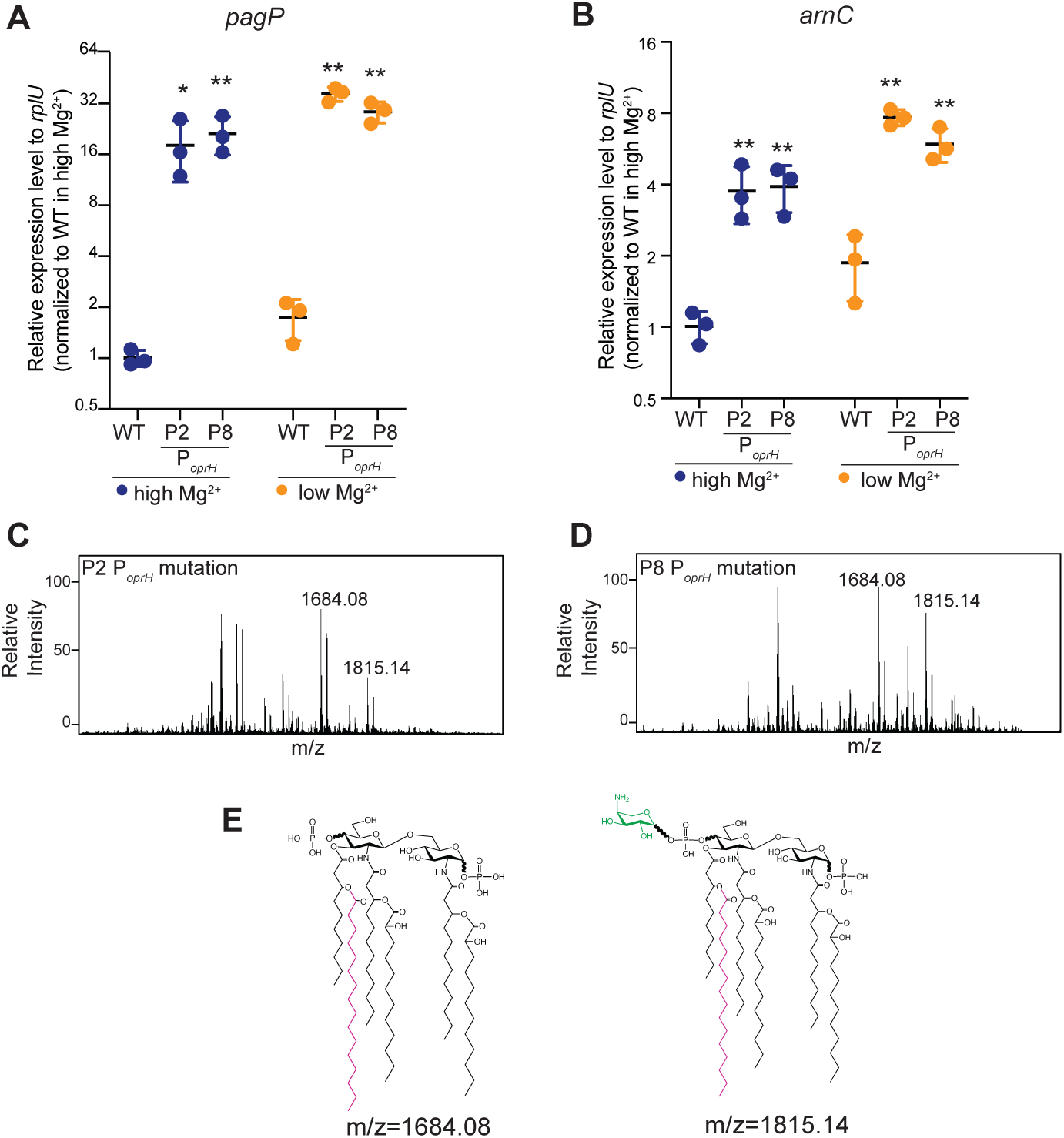
The PhoPQ pathway synergizes with early-arising mutations to confer colistin resistance. **(A-B)** Mutations in the promoter of the *oprH*/*phoP/phoQ* operon in replicate populations P2 and P8 cause the activation of PhoPQ-regulated genes *pagP* (A) and *arnC* (B), in both high and low Mg^2+^ conditions. (* *p* < 0.05, \*\**p* < 0.01, Dunnett’s one-way ANOVA test). **(C-D)** Mass spectra of lipid A modifications in PAO1 strain containing the promoter mutation in the *oprH*/*phoP/phoQ* operon from either P2 (C) or P8 (D) confirm the PhoPQ activation, with the *m/z* peak representing PhoPQ activity labeled in bold, where 1684.08 represents lipid A with PagP-mediated acylation but without L-Ara4N addition, and 1815.14 represents lipid A with PagP-mediated acylation and L-Ara4N addition. **(E)** Molecular structures of lipid A at m/z=1684.08, 1815.14 with PagP-mediated acylation (in magenta) and L-Ara4N addition (in green).

The identification of *phoPQ* mutations in two out of three evolutionary lines, compared to none in the *pmrAB* operon (Fig. 1B), suggests that these two systems played distinct roles in the development of colistin resistance during experimental evolution in low Mg^2+^. To test this possibility, we deleted the genes encoding the transcriptional factors PhoP and PmrA, individually or in combination. We first tested the effects of these mutations in the triple-mutant early backgrounds of P2, P5, and P8. The *phoP* deletion significantly reduced the colistin MIC to 3 μg/ml, whereas the *pmrA* mutation had only a mild effect. Moreover, *phoP pmrA* double deletion mutants showed no further reduction in resistance (Fig. S11A). This suggests that the PhoPQ, but not the PmrAB system, plays a dominant role in driving colistin resistance in the early stages of acquiring increased colistin resistance under low Mg^2+^ conditions. Our analysis of endpoint clones from the P2, P5, and P8 populations corroborated these findings; their high colistin resistance was much more dependent on PhoPQ than PmrAB (Fig. S11B). Our findings suggest that low Mg^2+^-triggered PhoPQ activation potentiated entirely novel evolutionary trajectories of colistin resistance that cannot be explored in high Mg^2+^ conditions.

### Two of three evolved lineages acquired colistin resistance by reducing colistin binding, but have compromised membrane integrity

We next investigated the potential cellular structural effects of the unique lipid A modifications by analyzing cell morphology with scanning electron microscopy (SEM). Our analyses revealed that endpoint clones from all three evolutionary lineages exhibited abnormal morphology compared to wild-type (WT) cells. These abnormalities were more severe in the P2 and P5 endpoint clones, which exhibited deformed membranes in both high and low Mg^2+^ conditions, with P5 cells being elongated in the low Mg^2+^ condition. In contrast, P8 endpoint clones exhibited less severe abnormalities, with slightly shorter cells than wild-type (WT) in both conditions (Fig. 5A and S12). Our findings suggested that cell morphology was affected in all three trajectories of cells that acquired high colistin resistance in low Mg^2+,^ but to variable degrees.

**Figure 5.**
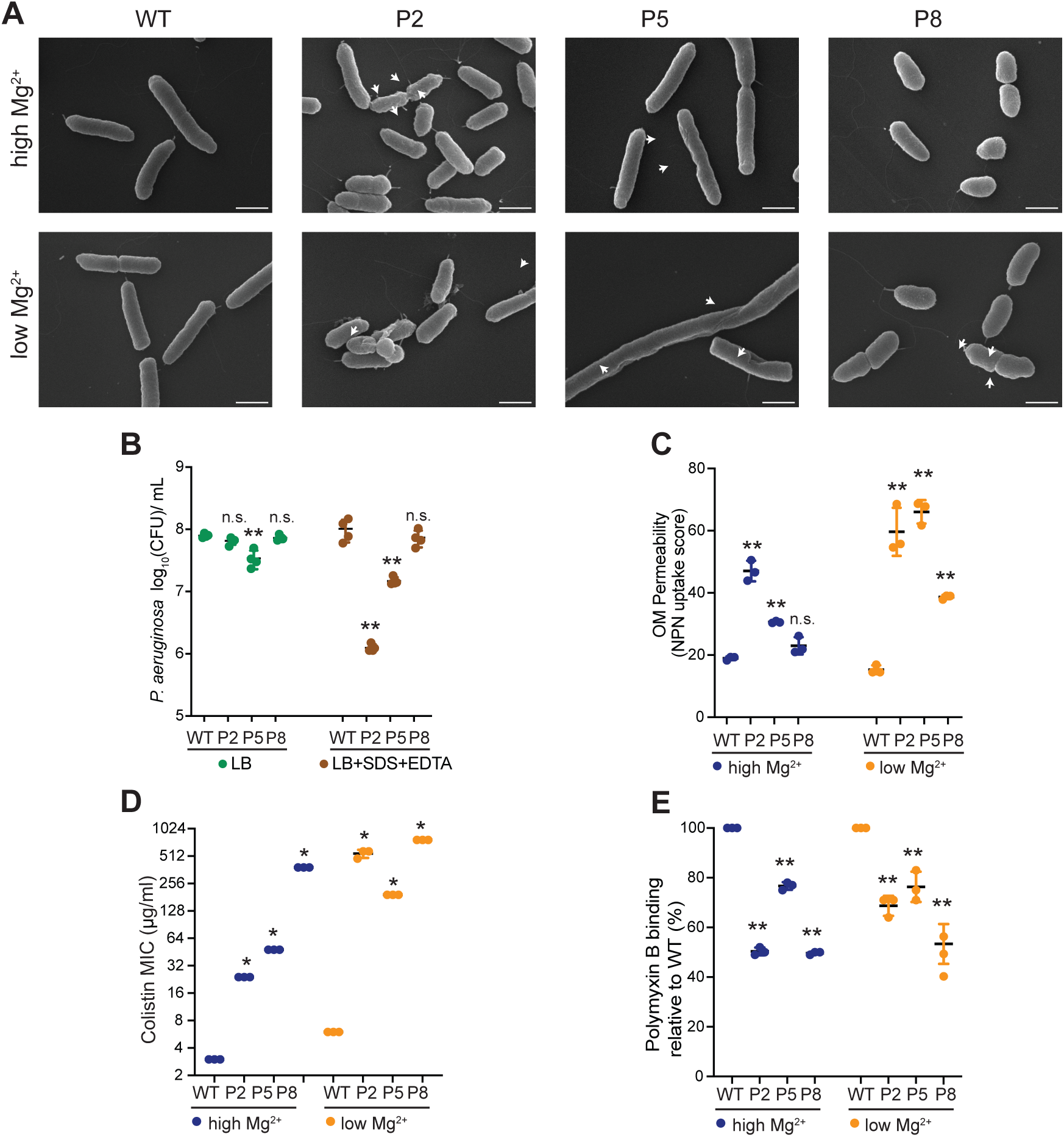
Replicate populations P2 and P5, but not P8, have membrane defects. **(A)** Scanning electron microscopic images of P2, P5, and P8 endpoint strains in high and low Mg^2+^ media. In high Mg^2+^, P2 and P5, but not P8, have discernible dents or kinks in the cell membrane (white arrows). In low Mg^2+^, P5 showed severe membrane deformation. All three displayed altered cell shapes compared to WT PAO1. The scale bar indicates 1μm. **(B)** Log-phase cells were serially diluted on an LB plate (green) or an LB plate supplemented with SDS and EDTA (brown) to assay bacterial resistance to membrane-perturbing agents. Cells from the P2 and P5, but not P8 lineages, were more sensitive to membrane stress. Mean ± std of 4 biological replicates is shown. (\*\**p* < 0.01, Dunnett’s one-way ANOVA test)**. (C)** An NPN assay was used to measure outer membrane permeability of WT, P2, P5, and P8. All three lineages had increased outer membrane permeability in the low-Mg^2+^ media. Mean ± std of 3 biological replicates is shown. (\*\**p* < 0.01, Dunnett’s one-way ANOVA test)**. (D)** Colistin MIC of endpoint clones in high and low Mg^2+^ media (* p<0.05, Mann-Whitney U test). **(E)** Dansyl-polymyxin B was used to quantify polymyxin B binding to WT PAO1 and endpoint clones in high and low Mg^2+^ media. All three endpoint clones had less binding to polymyxin B than WT in both conditions. Mean ± std of 3 biological replicates is shown. (\*\**p* < 0.01, Dunnett’s one-way ANOVA test)

Inspired by the SEM analyses, we queried whether the mutations that led to increased colistin resistance might have compromised outer bacterial membrane integrity. In a first assay, we grew endpoint clones in LB media and found that P2 and P8 endpoint clones exhibited similar viability to WT in LB alone, whereas P5 cells were growth-impaired even in LB alone. We then exposed the endpoint clones to membrane-perturbing agents such as SDS and EDTA [51] in LB media (Fig. 5B). Upon treatment with these agents, we found that P2 and P5 endpoint clones were significantly more sensitive than WT cells, while P8 cells remained relatively unaffected (Fig. S13). In a second assay, we assessed outer membrane permeability using the fluorometric probe NPN, which fluoresces upon entering the periplasm and binding phospholipids when membrane integrity is compromised in both high and low Mg^2+^ media. In high Mg^2+^, we found that P2 and P5 endpoint clones exhibited increased NPN uptake, whereas P8 did not (Fig. 5C). However, in low Mg^2+^ media, all three endpoint clones demonstrated higher NPN uptake than WT. Based on these findings, we conclude that P2 and P5 lineages have more severe membrane defects than the P8 lineage or the WT strain.

Although P2 and P5 endpoint clones have more permeable membranes, they exhibited greater resistance to polymyxin antibiotics, including colistin (polymyxin E) (Fig. 5D), and polymyxin B (Fig. S13A) than WT cells. To investigate how membrane-compromised cells gain increased resistance to antibiotics that target the outer membrane, we used dansyl-labeled polymyxin B [51] to quantify the binding of polymyxins to *P. aeruginos*a; dansyl-labeled polymyxin B fluoresces upon binding the hydrophobic portion of bacterial membranes. We used polymyxin B binding as a surrogate for how bacterial cells bind to all polymyxin antibiotics, including colistin. We validated the assay using Δ*phoP* Δ*pmrA* mutant cells and WT cells, finding that mutant cells minimally bind polymyxin B (Fig. S13B). Next, we evaluated polymyxin B binding by P2, P5, and P8 endpoint clones. In low Mg^2+^ media, we found that P2 and P5 had a 20-30% reduction in polymyxin B binding compared to WT, while P8, the most colistin-resistant clone, had the lowest levels of polymyxin B binding (Fig. 5E). Similar trends were observed in the high Mg^2+^ media (Fig. 5E). Our findings show that P2, P5, and P8 endpoint clones acquire a high degree of colistin resistance by reducing colistin binding. Two of these lineages do so through lipid A changes that compromise their membrane integrity—a novel mechanism of colistin resistance.

### Two modes of high colistin resistance lead to distinct patterns of fitness tradeoffs

We compared the relative fitness of all three endpoint clones to that of WT PAO1 in high or low Mg^2+^ media, in the absence of colistin. We found that the severity of their membrane defects correlated with fitness costs for the three evolved populations. P2 and P5 endpoint clones had significantly lower fitness than WT in both conditions, whereas P8 endpoint clones did not (Fig. 6A). Even the early triple-mutant reconstructed strains of P2 and P5 exhibited a significant fitness reduction compared to WT (Fig. S14), suggesting that a tradeoff between increased colistin resistance and bacterial membrane integrity manifests early during evolution in low Mg^2+^ conditions. Moreover, triple mutants and endpoint clones of these two populations had much lower colistin resistance in high Mg^2+^ media (Fig. S15A-B), consistent with our previous findings that their colistin resistance depends on PhoPQ activation. These findings explain why the P2 and P5 trajectory mutations do not (and likely cannot) occur under high Mg^2+^ conditions.

**Figure 6.**
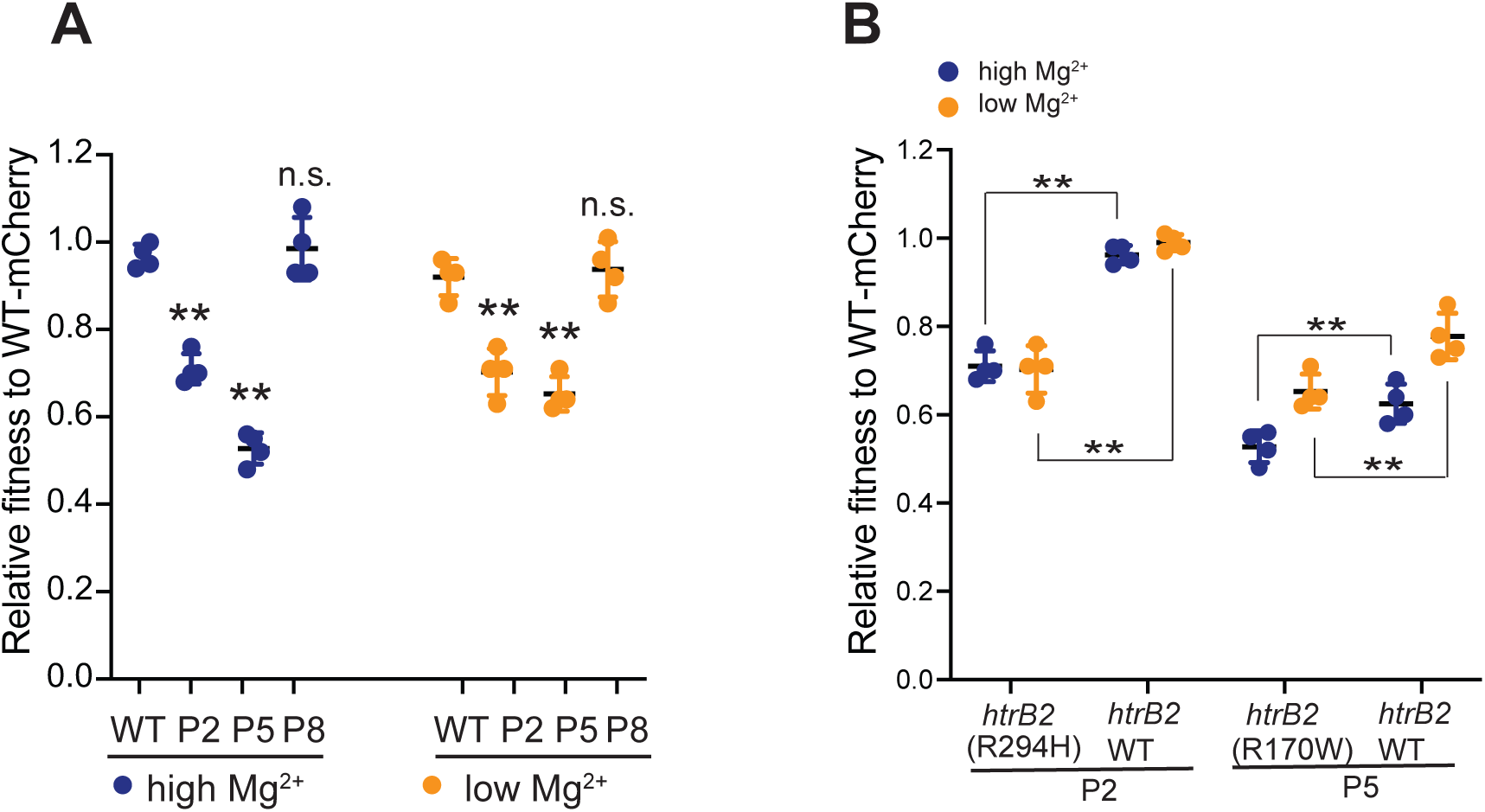
Colistin-resistant endpoint clones of P2 and P5 show fitness costs due to *hrtB2* mutations. **(A)** P2 and P5, but not P8, showed reduced fitness in high and low Mg^2+^. A competitive fitness assay was used to measure the fitness of P2, P5, and P8, relative to WT PAO1. Mean ± std of 3 biological replicates is shown. (\*\**p* < 0.01, Dunnett’s one-way ANOVA test) **(B)** *htrB2* reversion increases the fitness of P2 and P5 in high and low Mg^2+^ conditions. A competitive fitness assay was used to assess the fitness of each revertant strain relative to WT in high Mg^2+^ (blue) and low Mg^2+^ (orange) conditions without colistin. The Mean ± std of 4 biological replicates is shown. (\*\**p* < 0.01, \**p* < 0.05, one-tailed Mann-Whitney U test)

Based on mutations common to P2 and P5 but not found in P8 (Fig. 1B), we hypothesized that the loss-of-function and missense mutations in *htrB2*, an enzyme involved in lipid A acylation [47], were primarily responsible for membrane defects in P2 and P5. We tested this hypothesis by reverting the *htrB2* mutation to the WT allele in P2 and P5 endpoint clones. In both cases, we found that the *htrB2* reversion significantly improved bacterial fitness in low and high Mg^2+^ media (Fig. 6B). Additionally, an Δ*htrB2* mutant in WT PAO1 lowered membrane integrity [47], further confirming that *htrB2* mutations enhance colistin resistance at the expense of membrane integrity and bacterial fitness.

In contrast to P2 and P5, the P8 lineage followed a distinct evolutionary path, carrying a mutation in *lpxA,* the first essential gene in the biosynthesis of lipid A. Unlike P2 and P5, P8 endpoint clones showed only minimal impairment in bacterial membrane integrity and exhibited no fitness costs compared to WT cells, suggesting that this path incurs fewer tradeoffs. However, we found that the early P8 triple mutant also had significantly reduced fitness compared to WT in high, but not low, Mg^2+^ media (Fig. S14). This initial fitness cost may have been sufficient to prevent this trajectory from emerging under high Mg²⁺, whereas low Mg^2+^-dependent PhoPQ activation might have offset early fitness defects in P8. However, even this fitness cost in high Mg^2+^ conditions is ameliorated in P8 endpoint clones, suggesting that other late-occurring mutations (*PA5005*, *colS*, and *ftsY*) could suppress the previously acquired fitness defect.

### Dual modes of colistin resistance in low Mg^2+^ lead to distinct susceptibility to other antibiotics

Although membrane integrity defects lower colistin binding, thereby conferring colistin resistance, they may also confer susceptibility to other antibiotics that target intracellular processes, which the outer membrane typically protects against [52]. We hypothesized that the membrane integrity defects in P2 and P5 endpoint might increase susceptibility to other antibiotics by lowering the barrier to their penetration. To test this possibility, we examined the susceptibility of the P2, P5, and P8 end-point clones under high- and low-Mg^2+^ conditions to three antibiotics that are typically ineffective against *P. aeruginosa*: vancomycin, which targets the cell wall [53]; rifampicin, which targets RNA polymerase [54]; and azithromycin, which inhibits protein synthesis [55] (Fig. 7A).

**Figure 7.**
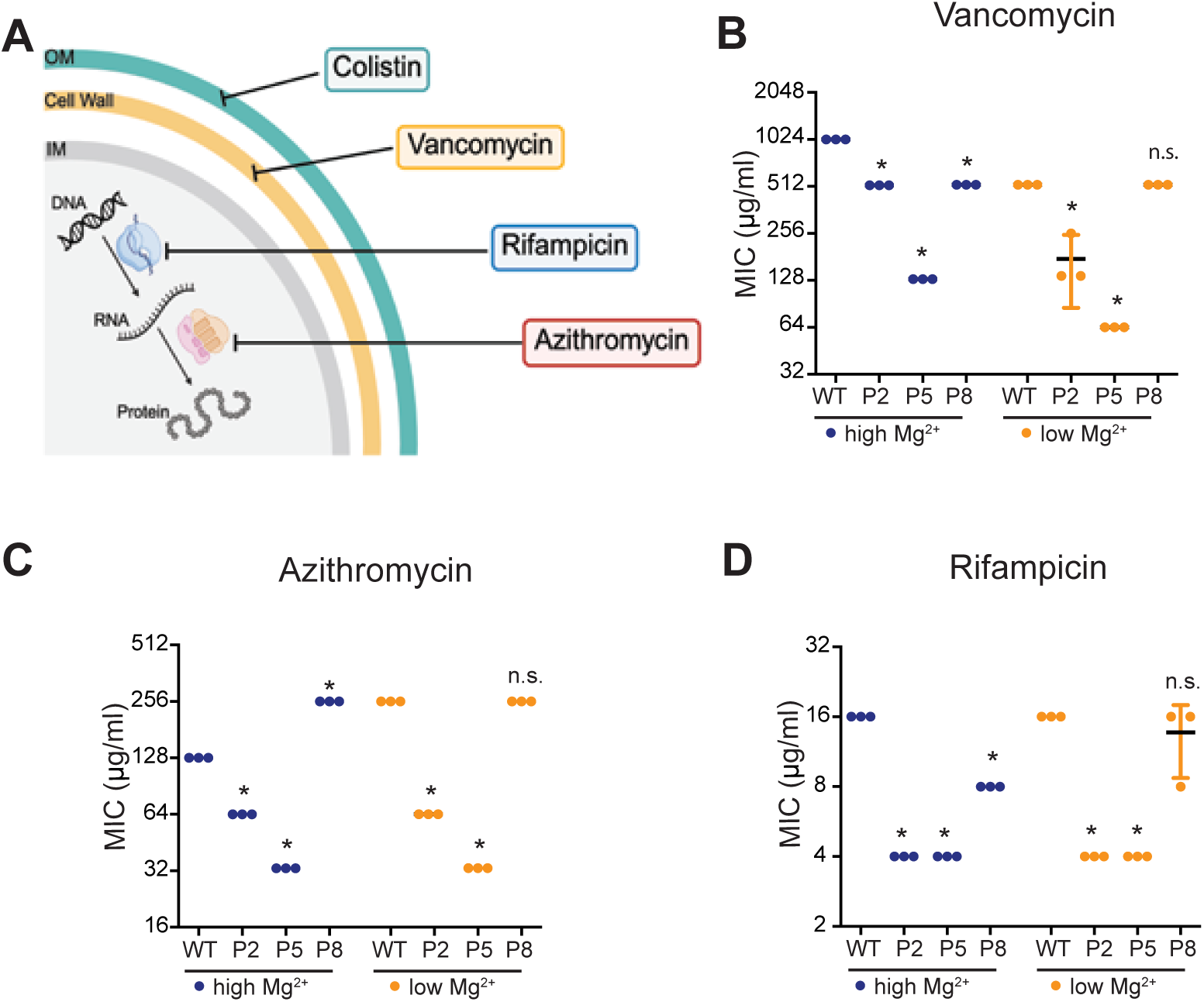
Colistin-resistant endpoint clones of P2 and P5 are more susceptible to other antibiotics. **(A)** A schematic of the mode of action of different antibiotics. **(B-D)** The resistance of endpoint clones to different antibiotics in high and low Mg^2+^ was measured using standard MIC assays with three biological replicates. MICs to vancomycin **(B)**, azithromycin **(C)**, and rifampicin **(D)** are shown. Mean ± std of 3 biological replicates is shown. (* p<0.05, Mann-Whitney U test)

As expected, the P2, P5, and P8 end-point clones exhibited much higher colistin resistance than WT in the low Mg^2+^ media (Fig. 5C). However, their susceptibility to other antibiotics correlated with membrane defects. P2 and P5 endpoint clones, which had *htrB2* mutations and more severe membrane defects, exhibited significantly lower minimum inhibitory concentrations (MICs) than wild-type (WT) cells for all three antibiotics (Fig. 7B-7D). By contrast, P8 endpoint clones, which have wild-type *htrB2* and only incur mild membrane defects, exhibited minimal increases in susceptibility to other antibiotics (Fig. 7B-7D). Our studies thus reveal two evolutionary paths of colistin resistance acquisition by *P. aeruginosa* that are only possible in low Mg²⁺ conditions. The genetic and biochemical underpinnings of these paths are distinct, as are their ensuing fitness tradeoffs.

## Discussion

Environmental conditions and microbial interactions with cohabiting species can strongly influence the evolution of microbial traits, such as antibiotic resistance [56–58]. Our study investigated how fungal-driven Mg^2+^ depletion influences the evolutionary pathways to colistin resistance in *P. aeruginosa* [37]. We identified convergent mutations in lipid A biosynthesis and modification genes, the *phoPQ* operon promoter, and *PA4824* (a putative Mg^2+^ transporter) that caused lipid A modifications and recapitulated early steps toward resistance under Mg^2+^-depleted conditions. This increased colistin resistance relied on synergistic interactions between the early mutations and PhoPQ activation under low Mg^2+^ conditions and came with fitness costs in high Mg^2+^ conditions, explaining why these mutations are not likely to occur in Mg^2+^-replete conditions.

Our analyses unexpectedly revealed two unique genetic and biochemical pathways to colistin resistance in low Mg^2+^ conditions. The first pathway (trajectories I and II, e.g. P2 and P5) relies on mutations in *htrB2,* which adds a laurate moiety to lipid A in the outer bacterial membrane of WT *P. aeruginosa* cells [47]. Examining the lipid A modifications in populations with *htrB2* mutations reveals that the missense mutations we identified are loss-of-function mutations, since no laurate modifications were detected (Fig. 3E-3F). As a result, populations with *htrB2* mutations have compromised outer membrane integrity that reduces colistin binding to the membrane. These genetic changes confer substantial fitness costs and heighten susceptibility to other antibiotics targeting intracellular processes. This finding represents a unique mode of the phenomenon of collateral sensitivity, in which increased resistance to one antibiotic leads to increased susceptibility to another [59, 60]. Interestingly, similar membrane vulnerabilities and increased antibiotic susceptibility have been reported in colistin-resistant *Acinetobacter baumannii* [61, 62], indicating that such LPS-mediated evolutionary tradeoffs may be widespread among gram-negative bacteria. Such tradeoffs support an evolution-guided strategy to combat antibiotic resistance by leveraging membrane defects to sensitize colistin-resistant bacteria to intracellular-targeting antibiotics.

The second pathway of low Mg^2+^-dependent colistin resistance relies on the synergy between PhoPQ activation and mutations in *PA4824* and *lpxA*, observed in trajectory III (e.g. P8). PhoPQ-dependent lipid A modification is known to enhance colistin resistance, but the potential role of LpxA in promoting resistance is a novel finding. LpxA encodes an essential acyltransferase that initiates the first step in the biosynthesis of lipid A [48], making loss-of-function mutations inviable. The *lpxA* mutation observed in our experiment does not appear to be a loss-of-function mutation. It might alter overall lipid A abundance rather than composition, which our biochemical analysis cannot detect, or have lipid A-independent function in promoting resistance. Moreover, this *htrB2*-independent pathway to colistin resistance does not significantly compromise outer membrane integrity, avoiding fitness costs and increased susceptibility to other antibiotics.

Although *htrB2*-dependent lineages exhibit fitness costs, their high frequency across replicate populations (6 out of 8), along with associated membrane defects, appears paradoxical. This pattern, however, may reflect the larger mutational target presented by *htrB2* compared to essential lipid A biosynthesis genes such as *lpxA*. Among the six *htrB2*-dependent populations, we identified three distinct mutations: R170W (P1 and P5), R294H (P2 and P3), and a 35 bp deletion (P7). In contrast, the two *lpxA*-mutant populations (P4 and P8) in trajectory III share the same G170C mutation. These findings suggest that *htrB2*-mediated resistance may arise more readily through multiple available mutational paths. Furthermore, the *htrB2*-dependent mechanism may involve a specific lipid A alteration that reduces colistin binding to the outer membrane, despite compromising membrane integrity.

In addition to *htrB2* and *lpxA*, one of the most intriguing genes identified in our evolution experiment is *PA4824*, a putative Mg^2+^ transporter [37]. All replicate populations incurred either the P224L or P244A mutation in *PA4824.* In two of three evolutionary lineages, reverting the P224L mutation to WT reduced colistin resistance. Yet, this mutation alone does not impair intracellular Mg^2+^ levels, unlike the complete loss of *PA4824* (Fig. S16), suggesting that resistance is promoted independently of Mg^2+^ uptake. Given that *PA4824* encodes a transmembrane protein, we suspect this mutation may stabilize the compromised outer membrane caused by other resistance-associated mutations. Future studies could investigate how the *PA4824* mutation, either alone or in combination with other mutations, affects the bacterial outer membrane to enhance colistin resistance.

Lipid A modifications that reduce colistin binding are a well-known means by which Gram-negative bacteria can acquire resistance to this antibiotic. Several mechanisms have been described, including PmrAB-dependent modification, which results in the addition of phosphoethanolamine or galactosamine to lipid A [63–65] or complete loss of LPS [62] in *Acinetobacter baumannii*, or presence of plasmid-borne phosphoethanolamine transferase in *P. aeruginosa* [66] and *Escherichia coli* [67]. Our work discovered several novel lipid A structures that can contribute to colistin resistance in *P. aeruginosa* by reducing polymyxin binding. In addition to PhoPQ-dependent modifications, we identified loss-of-function mutations in *htrB2* and *lpxO2*, which alter acyl chain number and hydroxylation status of lipid A and are necessary for increased colistin resistance in low Mg^2+^. Interestingly, identical *htrB2* and *lpxO2* mutations have been identified in a few clinical *P. aeruginosa* isolates [68–70], while other *lpxO2* loss-of-function variants have been observed in several *P. aeruginosa* isolates obtained from patients with cystic fibrosis [41, 71]. Though not previously linked to colistin resistance, we propose that these otherwise deleterious mutations may have been selected in Mg^2+^-poor environments by reducing colistin binding to membranes. Such unexpected lipid A modifications could be markers of low Mg^2+^-dependent colistin resistance in clinical *P. aeruginosa* isolates.

Our work also highlights the distinct roles of PhoPQ and PmrAB systems in shaping the evolutionary paths to colistin resistance. Although both two-component systems are activated upon Mg^2+^ depletion to modify lipid A [24, 29], we showed that selection for colistin resistance under chronic Mg^2+^ depletion drives PhoPQ activation—but not PmrAB activation—to synergize with several lipid A-altering mutations and enhance colistin resistance during evolution. Despite the fact that these two regulatory systems co-regulate several genes, we hypothesize that specific PhoP-regulated genes [49], which PmrAB does not activate, engage in key epistatic interactions with the mutations we identified. Beyond resistance, our data also reveal the importance of PhoPQ in maintaining cell morphology in low Mg^2+^ conditions. These findings suggest that PhoQ-specific kinase inhibitors could be used alongside colistin to prevent low Mg^2+^-dependent colistin resistance [72].

Together, our findings underscore the critical role of Mg^2+^ in shaping the evolution of antibiotic resistance. We demonstrate that bacterial evolution in low Mg^2+^ environments, such as during fungal-bacterial interactions [37], and within polymicrobial biofilms [38, 73], can lead to unconventional evolutionary pathways to resistance against colistin (and possibly other antibiotics). These adaptations often, but not always, incur trade-offs that can be exploited for therapeutic gain. Our study expands the mechanistic understanding of colistin resistance and highlights the ecological relevance of metal ions in driving the evolution of antibiotic resistance. By identifying genetic markers and vulnerabilities of low Mg^2+^-dependent resistance, our work lays a foundation for developing evolution-guided strategies to combat multidrug-resistant *P. aeruginosa* and other Gram-negative bacteria. More broadly, since many other antibiotic mechanisms are similarly dependent on metal ions [74–76], our work suggests that nutritional competition for metal ions may alter initial antibiotic resistance and potentiate new evolutionary pathways of antibiotic resistance.

## Materials and Methods

### Fungal and bacterial strains used in this study

Bacterial and fungal strains used in this study are listed in Table S2. Bacterial strains in this study were derived from *P. aeruginosa* PAO1. Unless otherwise specified, all experiments were performed in Brain Heart Infusion Broth (BHI, Sigma-Aldrich), buffered with 10% MOPS and 2% glucose to pH 7.0, followed by filter sterilization. All strains were grown at 30°C or 37°C. For antibiotics used in this study, 100 μg/mL gentamicin was used to select against *P. aeruginosa*. 50 μg/mL nystatin was used to select against *C. albicans*. Colistin (Sigma-Aldrich) was prepared as a 10 mg/mL stock solution.

### Co-culture colistin survival assay

All experiments began with bacterial or fungal cultures that were grown overnight. Starter cultures were diluted either 1:100 (for bacterial cultures) or 1:50 (for fungal cultures) in BHI and cultured for ∼4-5 hours to reach the logarithmic phase. Refreshed bacterial and fungal strains were added to BHI to reach the final bacterial cell density at 2.5 × 10^4^ cells/mL and fungal cell density at 5 × 10^5^ cells/mL for co-culture experiments. Identical amounts of bacterial or fungal cells were added separately in fresh BHI media as monoculture controls. These cultures were incubated for 18 h at 37°C with shaking. To equalize bacterial cell numbers in these two conditions before colistin treatment, the cell density of monoculture samples was adjusted to an optical density of 0.3 at 600 nm (OD_600_) to match the bacterial cell density after 18 h of growth in co-culture. Both equilibrated monoculture and co-culture samples were split into two 1 mL culture tubes. 192 µg/ml of colistin was added to one of the tubes. Cells in both tubes were incubated at 37°C for 1.5 hours, and the bacterial viability was determined by enumerating colony-forming units (CFU) in each condition. Bacterial survival was calculated as the ratio of CFU upon colistin treatment relative to no colistin.

### Antibiotic MIC assays

MICs (minimum inhibitory concentrations) were determined using a standard serial broth dilution method. Bacterial cells were cultured in BHI media overnight at 37°C with shaking and diluted 1:100 into 2 mL of either BHI media (high Mg^2+^ media) or *C. albicans*-spent media (low Mg^2+^ media) for 5 hours to determine MIC in high Mg^2+^ and low Mg^2+^ conditions, respectively. Next, 3 μL of the log-phase culture was inoculated into each well with 200 μL of the same media and titrated antibiotics (colistin from 0 to 768 μg/mL, azithromycin from 0 to 128 μg/mL, vancomycin from 0 to 1024 μg/mL, and rifampicin from 0 to 256 μg/mL). The plate was incubated at 37°C for 24 h, after which OD_600_ was measured using a microplate reader. The MIC of each strain was determined as the lowest concentration of antibiotics at which OD_600_ was half of the maximum OD_600_ observed in the absence of colistin. Three technical replicates were used to measure the MIC of each strain.

### Mutation reconstruction in WT PAO1

We used an allelic exchange method to reconstruct mutations in WT PAO1 or revert evolved mutations to a WT allele in endpoint clones [77]. Briefly, a gene fragment with or without mutation was incorporated into the pEXG2 plasmid and transformed into *E. coli* S17. The S17 donor was subsequently mixed with *P. aeruginosa* recipients on an LB agar plate at a 5:1 ratio of donor to recipient cells, and the cell mixture was incubated at 30°C overnight. The cell mixture was plated on LB agar plates with 100 μg/mL gentamicin to select for cells containing the deletion plasmid integrated into the *P. aeruginosa* genome. Counterselection was performed with LB agar plates containing sucrose. Gene deletions were confirmed by Sanger sequencing of PCR products (GeneWiz, Azenta). Primers and plasmids used for strain constructions are listed in Table S3.

### Whole genome sequencing of evolutionary intermediates of evolved populations

Populations were frozen at two-week intervals and at the endpoint of the experimental evolution [37]. We revived each population from a freezer stock and grew them for 24 h. For co-culture evolved populations, we treated them with nystatin to remove fungal cells. A 3 ml bacterial culture was collected for DNA extraction using the DNeasy Blood & Tissue kit (Qiagen). Sequencing libraries were made and commercially sequenced using Illumina technology by SeqCenter (https://www.seqcenter.com/). Variants were called using the breseq software v0.37.1, with the *P*. *aeruginosa* PAO1 genome (GCF_00006765.1) as the reference genome. The average genome coverage was 98.8 fold (+/− 0.04). Mutations were manually curated by identifying unique variants in each evolved population compared to the ancestor. Whole-genome sequencing data of evolved populations are available in NCBI Bioproject PRJNA1251133.

### Mass Spectrometry Analysis of lipid A

Lipid A structural analysis was performed using Fast Lipid Analysis Technique (FLAT), as previously described [78]. Bacterial cultures were grown to the mid-log phase and pelleted. The pellet was scraped directly onto a steel MALDI target plate in duplicate with a pipette tip. FLAT extraction buffer (1 µL; 0.2 M anhydrous citric acid, 0.1 M trisodium citrate dihydrate, pH 4.5) was pipetted over bacterial spots. The FLAT target plate was incubated at 100°C in a humidified heat block for 30 minutes. Bacterial spots were washed with endotoxin-free water Gibco (Grand Island, NY, USA) and air-dried, followed by the addition of norharmane matrix (10mg/mL in 1:2 MeOH : CHCl3 – Sigma Aldrich, St. Louis, MO, USA). Mass spectra were collected in negative-ion mode using a Bruker timsTOF Flex. Agilent ESI Tune Mix was used as an external calibrant. Mass spectra were processed using DataAnalysis v3.4 software. Ions for tandem mass spectrometry were identified in DataAnalysis and fragmented in the gas phase between 70 and 90 eV. Each MS analysis was performed with at least two independent biological replicates. Mass spectra were analyzed using mMass v5.5.0. Table S1 describes all the m/z values determined in our experiments.

### Quantification of gene expression

Log-phase cell cultures in BHI and *C. albicans*-spent BHI were collected for RNA extraction using the RNAeasy mini kit (Qiagen) following the commercial protocol. RNA was converted to cDNA using qScript cDNA synthesis kit (QuantaBio). The expression level of genes of interest was determined by quantitative PCR using PowerTrack SYBR PCR mix (ThermoFisher) and gene-specific primers (Table S3). Gene expression level was determined relative to a housekeeping gene, *rplU*.

### Outer membrane permeability assay

We used a previously established NPN uptake assay [18, 79]. Bacterial cells were grown to log phase in BHI or *C. albicans*-spent BHI. Cells were diluted to an OD_600_ 0.5 in 1ml 5mM HEPES, pH 7.2. 150 μl bacterial suspension was added to wells of a black microtiter plate with clear-bottomed wells. The fluorescent probe N-phenyl-1-naphthylamine (NPN, Sigma Aldrich) was added to a final concentration of 10 μM. Fluorescence was measured immediately in a Cytation 5 plate reader using an excitation wavelength of 535 nm and an emission wavelength of 405 nm. Fluorescence measurements were obtained every 1min for 15 mins, and the degree of outer membrane permeability, referred to as the NPN uptake score, was calculated using the following equation: *(Fluorescence of sample with NPN -Fluorescence of sample without NPN)/ (Fluorescence of HEPES buffer with NPN – Fluorescence of HEPES buffer without NPN)*.

### Scanning electron microscopy experiments

1mL log-phase cells in BHI were collected and fixed with formaldehyde. Duplicates of 50 µL of each sample were applied in a pool on poly-l-lysine coated coverslips for 30 min, rinsed with 0.1M sodium cacodylate buffer, and treated with 1% osmium tetroxide for 1 hour. The coverslips were then rinsed with cacodylate buffer, dehydrated through a graded series of alcohols, infiltrated with HMDS (Electron Microscopy Sciences, Hatfield, PA), and allowed to air dry. Coverslips were mounted on stubs and sputter-coated with gold/palladium (Denton Desk IV, Denton Vacuum, Moorestown, NJ). Samples were imaged on a JSM 6610 LV scanning electron microscope at 15kV and 12mm working distance (JEOL, Tokyo, Japan).

### Polymyxin B binding assay

Bacterial strains were cultured in BHI media overnight at 37 °C with shaking. Cultures were then diluted at 1:100 in BHI (high Mg^2+^) and *C. albicans*-spent BHI (low Mg^2+^) for 5 hours to reach log phase. 1mL of log-phase cells were collected, washed, and resuspended in 1mL 0.9% NaCl to reach 0.5 OD_600_. 1mL of cell suspension was then mixed with 3 µg/mL dansyl-polymyxin B (Sigma-Aldrich) for 30 minutes in dark. After incubation, 150 µL of each suspension was transferred into wells of a black 96-well microtiter plate to measure fluorescence (excitation at 340 nm and emission at 485 nm). The dansyl fluorophore shows low fluorescence in aqueous solution but exhibits enhanced fluorescence when bound to LPS of bacterial membranes. To calculate polymyxin B binding to cells, we use the following formula to measure changes in fluorescence intensity: 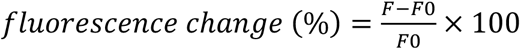, where *F* = fluorescence intensity of sample with dansyl-polymyxin B, and *F0* = fluorescence intensity of sample with dansyl-polymyxin B. The relative fluorescence intensity of each sample was normalized to WT under the same condition.

### Competitive fitness assay

WT PAO1 with chromosomally integrated mCherry was used as the reference strain in fitness competition with evolved strains. The relative fitness of each evolved strain was determined either in BHI media only (high Mg^2+^ condition) or in BHI media in co-culture with *C*. *albicans* (low Mg^2+^ condition) to recapitulate the culturing conditions of experimental evolution. All bacterial and yeast strains were incubated in BHI at 37°C to log phase. A non-fluorescent test strain was mixed with the fluorescence-labeled reference strain at a ratio of 1:1. Part of the cell mixture was used to determine the initial ratio of sample and reference strains using flow cytometry (BD FACSymphony A5 Cell Analyzer). Then, 10 μl of cell mixture was inoculated separately in 2 ml BHI alone (monoculture) or 2 ml BHI with 2 × 10^5^ *C*. *albicans* cells (co-culture) and grown in BHI at 37°C for 18 hours. After 18 hours, monoculture samples were diluted 100-fold to measure the ratio of sample and reference strain. For co-culture samples, a low-speed spin (500x*g* for 8 min) was applied to separate bacterial and fungal populations. Supernatants enriched with bacterial cells were identified using flow cytometry. At least 30,000 cells were collected for each sample, and the data were visualized using FlowJo 10.4.1. The reference strain was cultured separately to estimate the number of generations during the experiment. Each experiment was conducted in at least two biological and two technical replicates. To calculate the relative fitness, *w*, of each sample strain to the reference strain, we followed the formula: *w* = 1+*s*,

where *s* is selection coefficient: 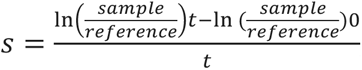,

where *t* = number of generations and (sample/reference) is the ratio between a sample strain and the reference strain [80].

## Acknowledgments

We thank the Ernst, Dandekar, and Malik lab members for their valuable discussions on this project. We thank Pete Greenberg, Carrie Harwood, Nina Salama, and Andrew Murray for their comments on the manuscript. We especially thank Nina Salama for her suggestion to investigate lipid A modifications and Steve MacFarlane and the Fred Hutch Electron Microscopy & CryoEM Core for supporting scanning electron microscopy experiments.

The following agencies funded this study:

Cystic Fibrosis Foundation HSIEH24F0 and HSIEH21F0-CI (to YPH)

Cystic Fibrosis Foundation ERNST23G0 and NIH AI104895 (to RKE)

NIH R35 GM152107 (to AAD)

Howard Hughes Medical Institute Investigator award (to HSM).

Funding agencies played no role in the study design or the decision to publish.

## Author contributions

Conceptualization: YPH, IPO, RKE, AAD, HSM

Methodology: YPH, IPO, ZW, WS

Investigation: YPH, IPO, ZW, WS, HY, LMV

Visualization: YPH, IPO, ZW, WS, AAD, HSM

Funding acquisition: YPH, RKE, AAD, HSM

Supervision: YPH, RKE, AAD, HSM

Writing-original draft: YPH, IPO

Writing-review & editing: WS, ZW, HY, LMV, RKE, AAD, HSM

## Data and materials availability

All data in the main text and supplementary materials are publicly available. Strains will be shared upon request.

