## supplementary figures and tables for "Magnesium depletion unleashes two unusual modes of colistin resistance with different fitness costs"

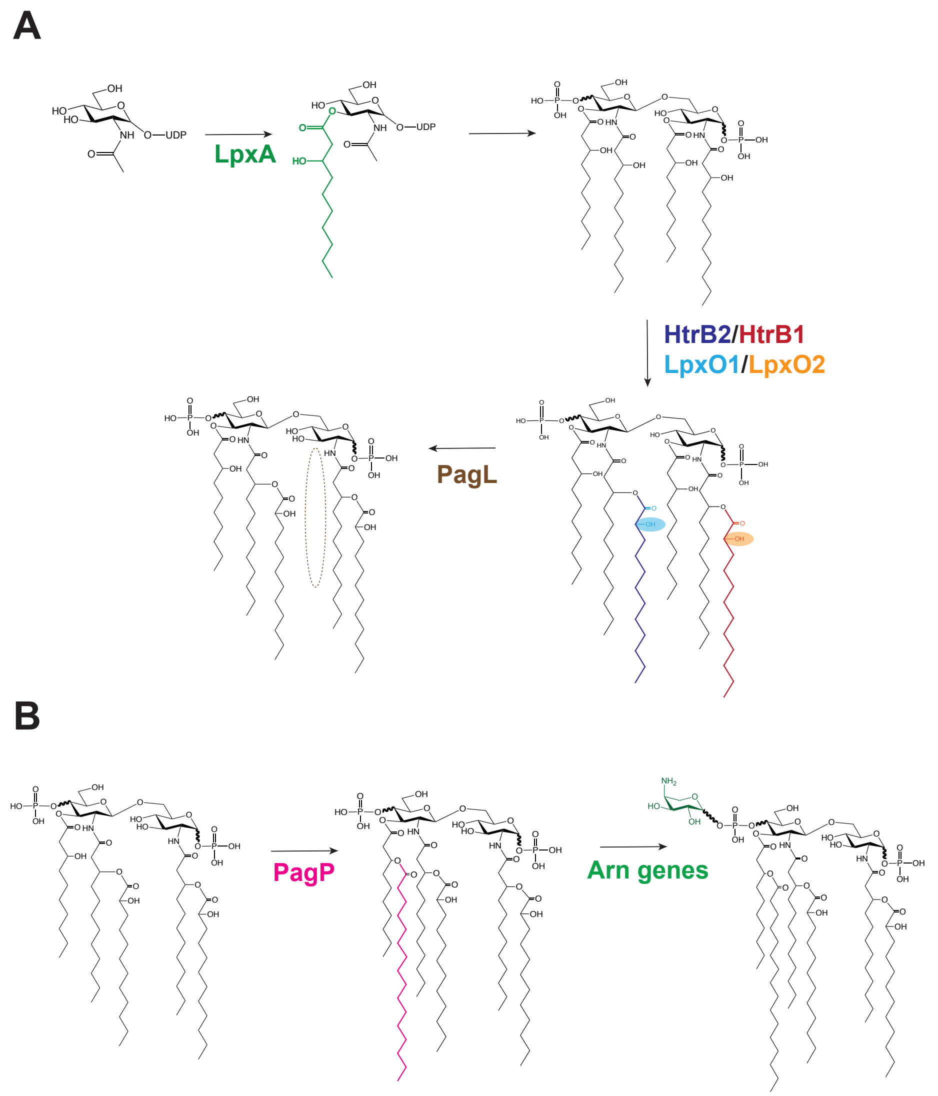


**Fig. S1 Schematic of lipid A biosynthesis and modification pathway**
**(A)** Lipid A biosynthesis in gram-negative bacteria begins with the synthesis of lipid $IV_A$, which consists of phosphorylated glucosamine sugars, each of which serves as the backbone for an N-linked 3'OH-C12 and O-linked 3'OH-C10 acyl chain in *P. aeruginosa*. First, LpxA catalyzes the transfer of an acyl group to UDP-N-acetylglucosamine (UDP-GlcNAc), forming UDP-3-O-acyl-GlcNAc. Subsequently, the HtrB1/LpxO2 and HtrB2/LpxO1 acyltransferases and hydroxylases add secondary O-linked C12 acyl chains and 2' hydroxyl groups at the 3'OH residue of the N-linked acyl chains; such regulation is very conserved in gram-negative bacteria. Lipid biosynthesis/ modification mutations observed in our evolution experiment are bolded (LpxA, HtrB2, and LpxO2). In certain cases, PagL removes the acyl chain from the 3-position of lipid A. **(B)** Low $Mg^{2+}$-responsive PhoPQ pathway upregulates enzymes that modify lipid A structures [29, 30]. These include *pagP*, which encodes an acyl transferase that adds a C16 fatty acid, and the *arn* operon (*arnBCADTEF*), which attaches 2-amino-2-hydroxy-L-arabinose (aminoarabinose) to the phosphate groups on the glucosamine backbone of lipid A.

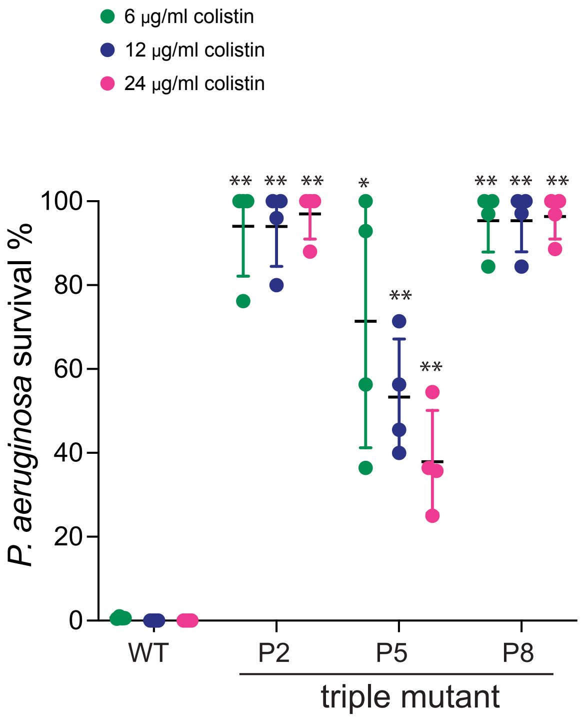


**Fig. S2 Triple-mutation-reconstructed strains show significantly increased survival to a range of low concentrations of colistin in co-culture**

*P. aeruginosa* survival to low concentrations of colistin in co-culture was measured using the colistin survival assay. Survival of triple mutants to 6, 12, and 24 µg/ml colistin is shown in green, blue, and magenta, respectively. P2-derived triple mutant contains *htrB2*, *PA4824*, and the *oprH*/*phoPQ* promoter mutations. P5-derived triple mutant contains *htrB2*, *PA4824*, and *lpxO2* mutations. P8-derived triple mutant contains *lpxA*, *PA4824*, and the *oprH*/*phoPQ* promoter mutations. All three triple mutants had higher survival to these concentrations of colistin, compared to WT. Mean ± std of 4 biological replicates is shown. The survival of WT PAO1 in three conditions is used as the reference for statistical analysis (**$p < 0.01$, *$p < 0.05$, Dunnett's one-way ANOVA test).

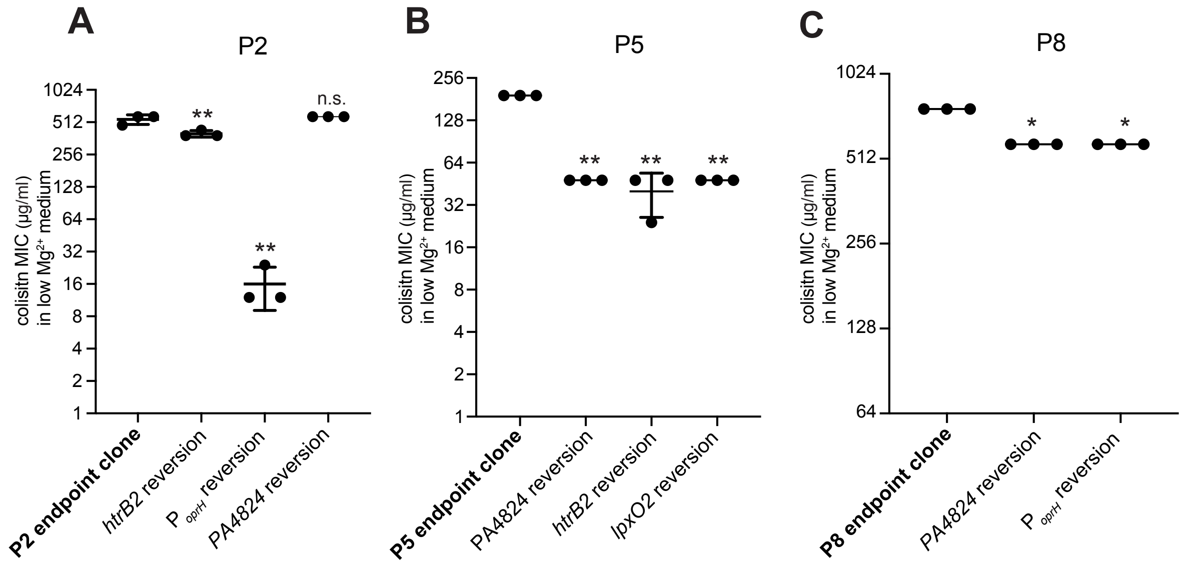


**Fig. S3 Mutation reversion strains show reduced colistin MIC in low $Mg^{2+}$ media**
Colistin resistance of endpoint clones and single-mutation reversion strains was measured by colistin MIC assays. Colistin resistance of strains derived from P2, P5, and P8 is shown in A, B, and C. Mean ± std of 3 biological replicates is shown and the MIC of each endpoint clones is used as the reference for statistical analysis (**$p < 0.01$, *$p < 0.05$, Mann-Whitney U test).

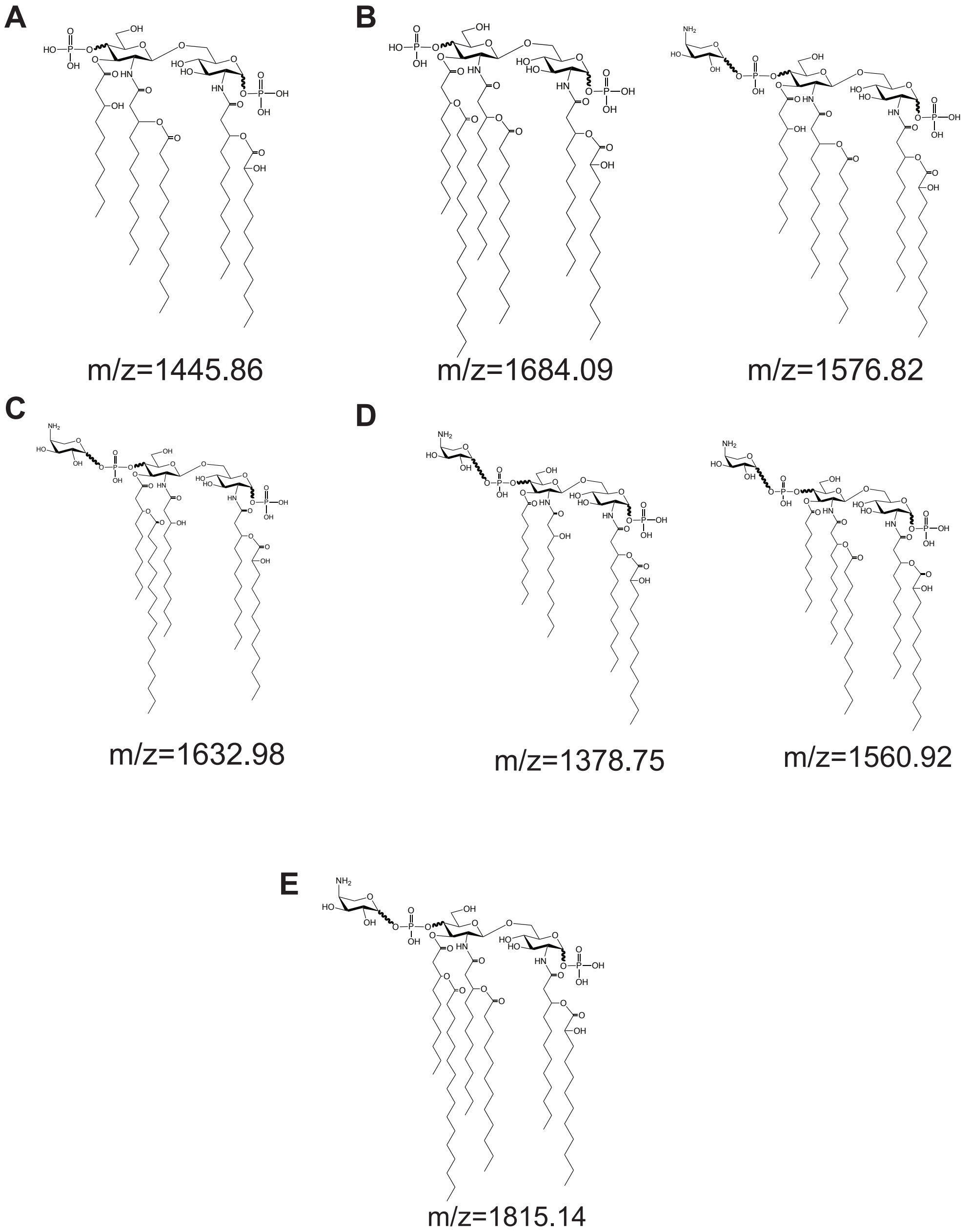


**Fig. S4 Molecular structures of lipid A in the WT PAO1 strains and endpoint clones**
**(A)** Molecular structure of lipid A (m/z=1445.86) of WT PAO1 in high $Mg^{2+}$ media. **(B)** Molecular structure of lipid A (m/z=1576.82) of WT PAO1 in low $Mg^{2+}$ media. **(C)** Molecular structure of lipid A (m/z=1362.98) of P2 in low and high $Mg^{2+}$ media. **(D)** Molecular structures of lipid A (m/z=1378.75, 1560.92) of P5 in low and high $Mg^{2+}$ media. **(E)** Molecular structure of lipid A (m/z=1815.14) of P8 in low and high $Mg^{2+}$ media.

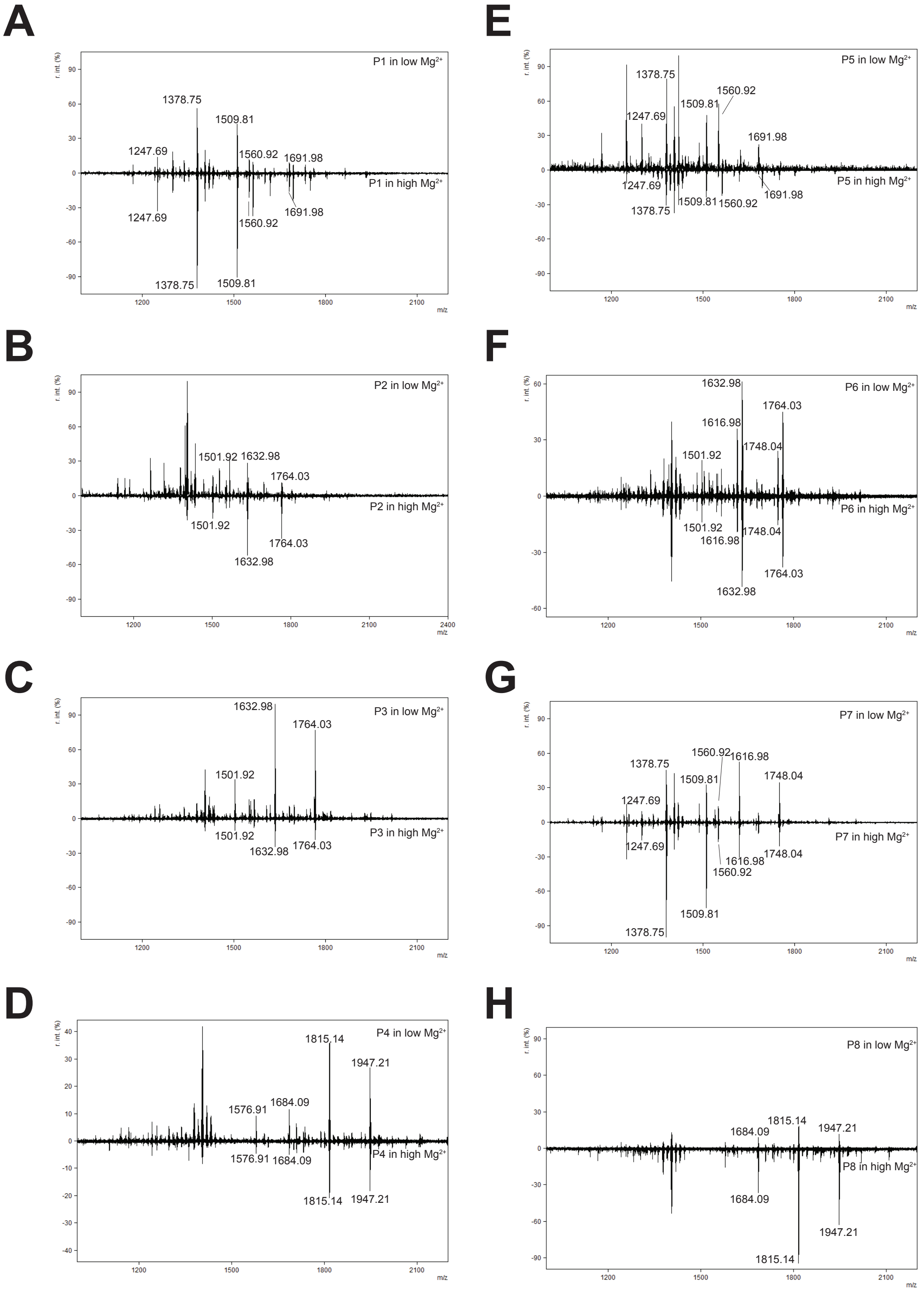


**Fig. S5 Mass spectra of evolved populations in high and low $Mg^{2+}$ conditions are comparable**

**(A-H)** Mass spectra of lipid A for P1 through P8 are shown in low (top) and high $Mg^{2+}$ (bottom) conditions. All lipid A peaks present in low $Mg^{2+}$ media are also present in high $Mg^{2+}$ conditions. Variations in total peak height are due to variations in absolute peak intensity between samples. Populations 1, 5, and 7 contain tetra-acylated lipid A lacking HtrB2-mediated C12 addition, penta-acylated lipid A with HtrB2-mediated C12 addition, and all lipid A lacks LpxO1/2-mediated 2’-hydroxylation. Populations 2, 3, and 6 contain penta-acylated lipid A with PagP-mediated palmytoylation and a lack of HtrB2-mediated C12 addition. Populations 4 and 8 contain hexa-acylated lipid A with PagP-mediated palmytoylation, and are singly 2’-hydroxylated, lacking LpxO1-mediated 2’-hydroxylation. See Table S1 for further details of every lipid A structure.

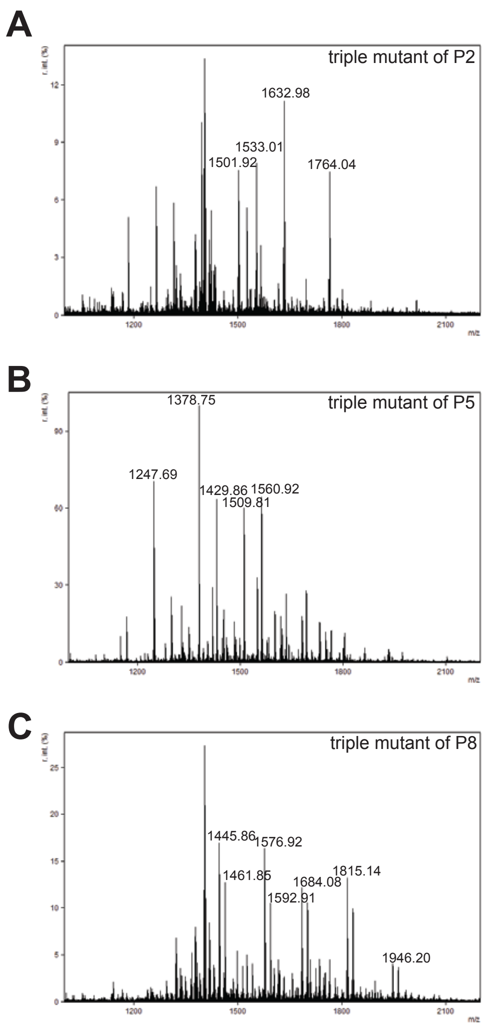


**Fig. S6 Mass spectra of triple-mutation-reconstructed strains are similar to those of endpoint populations**

Mass spectra of lipid A from all triple mutants. **(A)** Lipid A MS of P2 triple reconstruction mutant. Peaks at m/z 1501.92, 1632.98, and 1764.04 were present in the P2 endpoint clone, representing penta-acylated lipid A with PagP-mediated C16 addition and no HtrB2-mediated C12 addition. **(B)** Lipid A MS of the P5 triple mutant reconstruction. All peaks present were also present in the P5 endpoint clone, with peaks at m/z 1247.69, 1378.75, and 1509.81 representing tetra-acylated lipid A lacking HtrB2-mediated C12 addition. Peaks at m/z 1429.86 and 1560.92 represent penta-acylated lipid A containing HtrB2-mediated C12 addition, but no LpxO1 or LpxO2-mediated 2'-hydroxylation. **(C)** Lipid A MS of P8 triple mutant reconstruction. Peaks at 1684.08, 1815.14, and 1946.20 were all present in the P8 endpoint clone, representing hexa-acylated lipid A with a single 2'-hydroxylation mediated by LpxO2, and PagP-mediated palmytoylation. Peaks at 1445.86 and 1576.92 were not present in the P8 endpoint clone and represent penta-acylated lipid A without PagP-mediated palmitoylation. All peaks that are +15.99 m/z from previously mentioned peaks contain LpxO1-mediated 2-hydroxylation, which is also not present in the original P8. See Table S1 for further information of all lipid A structures present.

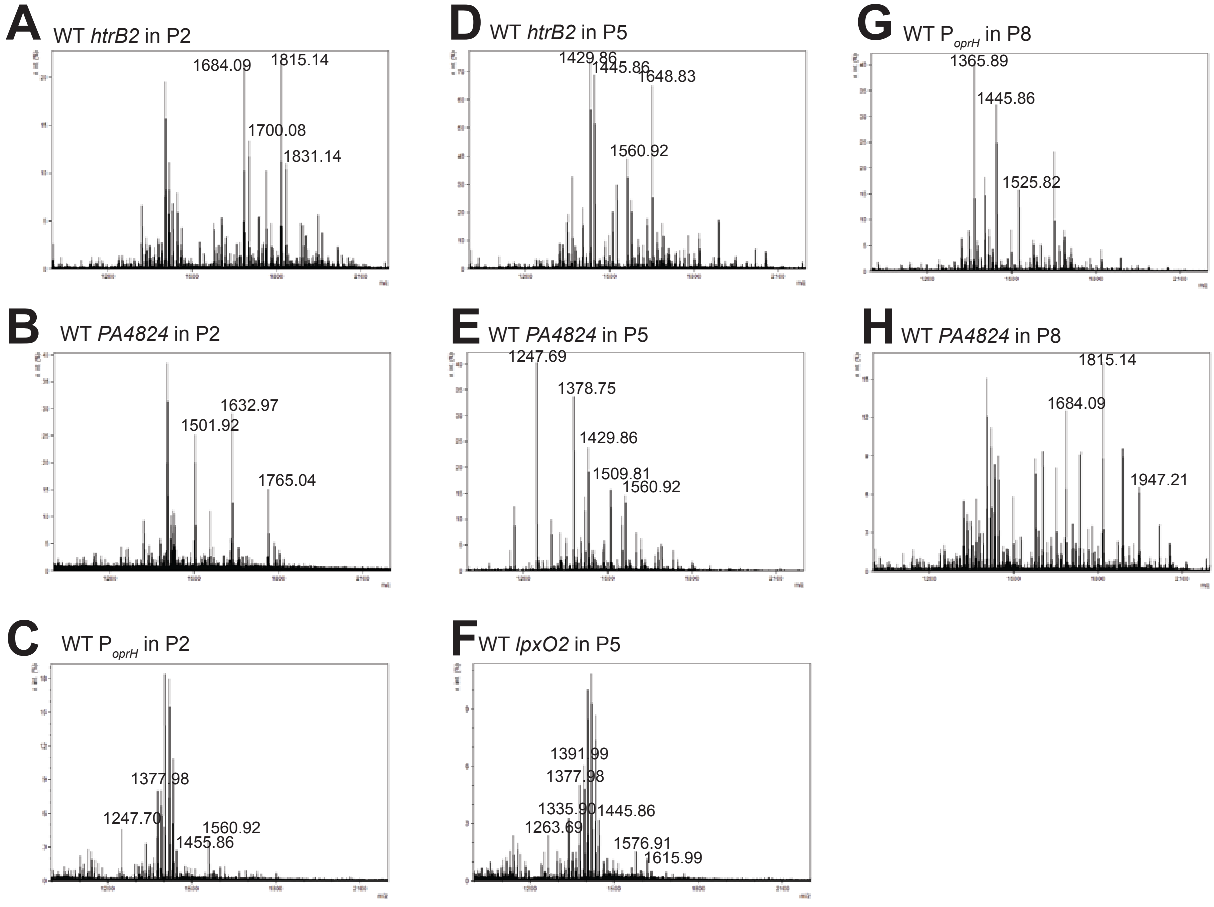


**Fig. S7 Mass spectra of mutation reversion strains**

Mass spectra of lipid A from all mutation reversion strains. **(A)** MS of WT *htrB2* in P2. Peaks present indicate hexa-acylated lipid A with HtrB2 function restored. **(B)** MS of WT *PA4824* in P2. All peaks present were also found in the original P2. **(C)** WT o*prH*/*phoPQ* promoter in P2. Peaks present indicate tetra- and penta-acylated lipid A lacking in palmitoylation. Single aminoarabinose addition peaks are still present. **(D)** MS of WT *htrB2* in P5. Peaks present indicate penta-acylated lipid A with the restored function of htrB2. **(E)** MS of WT *PA4824* in P5. All peaks present were also found in the original P5. **(F)** WT *lpxO2* in P5. Peaks present include tetra- and penta-acylated, singly hydroxylated lipid A. **(G)** MS of WT *oprH*/p*hoPQ* promoter in P8. Peaks present are penta-acylated lipid A varying in phosphorylation level, indicating a lack of PagP-mediated palmitoylation, as well as a lack of L-Ara4N addition. **(H)** MS of WT *PA4824* in P8. All peaks present were also found in the original P8. See Table S1 for further details of all lipid A structures present.

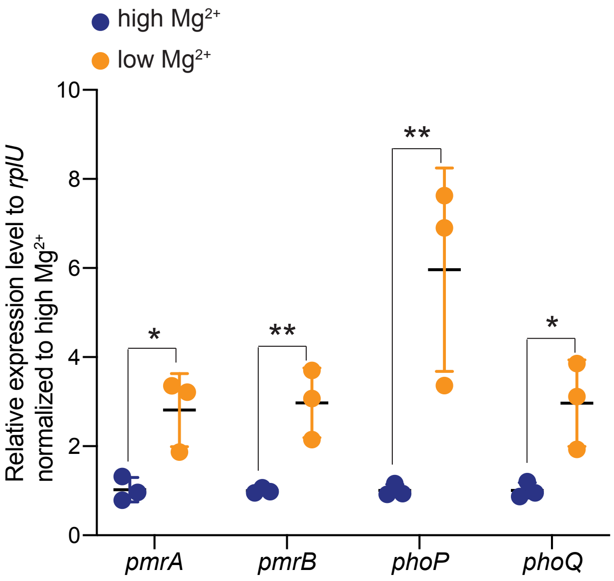


**Fig. S8 *phoP, phoQ, pmrA, and pmrB* genes were induced when WT PAO1 grew in low $Mg^{2+}$ media**

Log-phase cultures of the WT PAO1 in high and low $Mg^{2+}$ media were processed for RNA extraction. QPCR with gene-specific primers was used to determine the expression of *phoP, phoQ, pmrA,* and *pmrB* genes, as described in the Materials and Methods. Gene expression is shown as the expression levels relative to the internal control rplU and normalized to high $Mg^{2+}$ media. All these genes were upregulated in the low $Mg^{2+}$ media. Mean ± std of 3 biological replicates are shown (**$p < 0.01$, *$p < 0.05$, one-tailed Mann-Whitney U test).

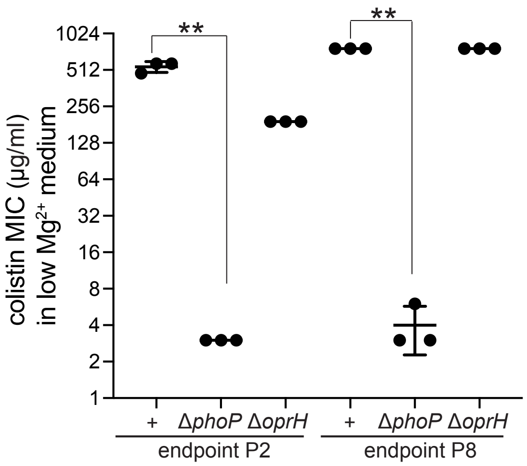


**Fig. S9 Deletion of *phoP*, but not *oprH*, in P2 and P8 clones reduced colistin resistance.** *phoP* or *oprH* was deleted in the P2 and P8 endpoint clones to distinguish which gene is required for evolved colistin resistance. Colistin resistance was measured by MIC in the low $Mg^{2+}$ media. *phoP* deletion, but not *oprH* deletion, significantly reduced colistin MIC. Mean ± std of 3 biological replicates is shown. (**p* < 0.05, One-tailed Mann-Whitney U test)

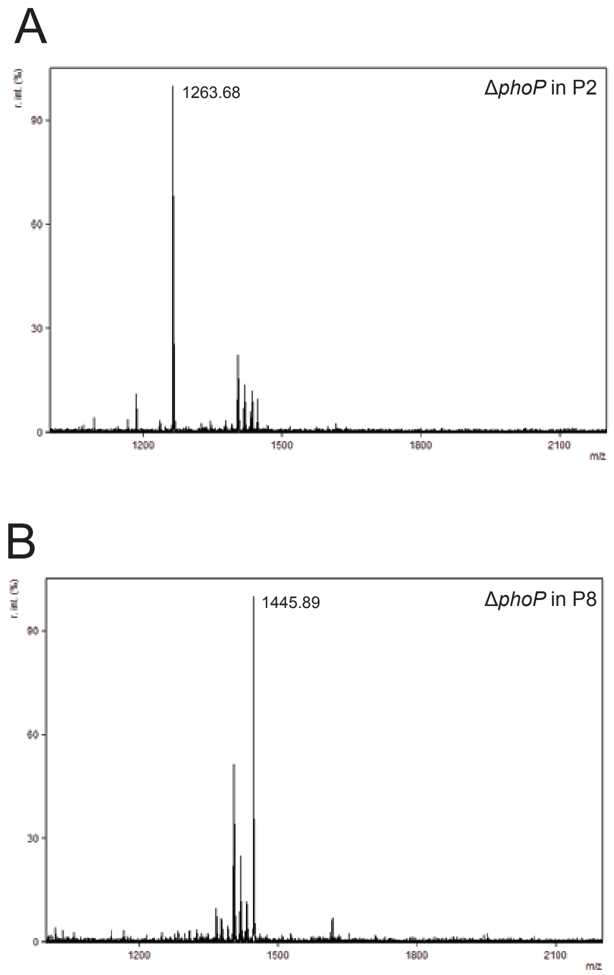


**Fig. S10 Mass spectra of Δ*phoP* in the endpoint strains of P2 and P8**

Δ*phoP* in P2 (A) and P8 (B) show lipid A without aminoarabinose addition and PagP-mediated acylation. (A) Mass spectra of Δ*phoP* in P2 yielded only one lipid A peak at 1263.68 *m/z*. This corresponds to a tetra-acylated lipid A structure lacking the previously present aminoarabinose additions and PagP-mediated palmytoylation in P2. (B) FLAT followed by MALDI-TOF MS of Δ*phoP* in P8 yielded only one major lipid A ion corresponding to the penta-acylated, singly hydroxylated 1445.89 *m/z*, lacking the aminoarabinose additions and PagP-mediated C16 addition that were present in the final P8 evolved clone.

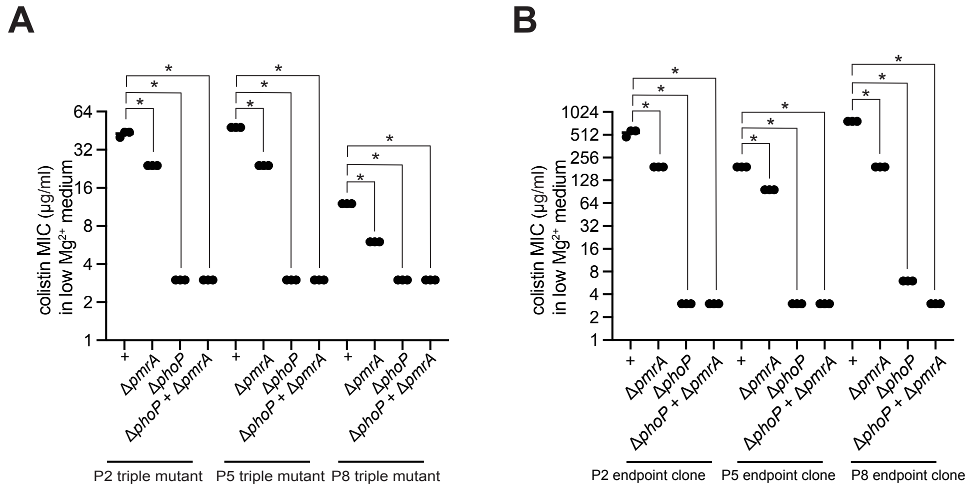


**Fig. S11 Colistin resistance in P2, P5, and P8 requires PhoPQ activity.**
Δ*pmrA*, Δ*phoP*, and Δ*phoP* Δ*pmrA* were made in the triple mutant **(A)** and endpoint clones **(B)**. Colistin resistance was measured by the standard MIC assay in the low $Mg^{2+}$ media. In both evolutionary backgrounds, *phoP* deletion significantly reduced colistin MIC, but *pmrA* deletion had a mild effect. Mean± std of 3 biological replicates are shown. (*$p < 0.05$, One-tailed Mann-Whitney U test)

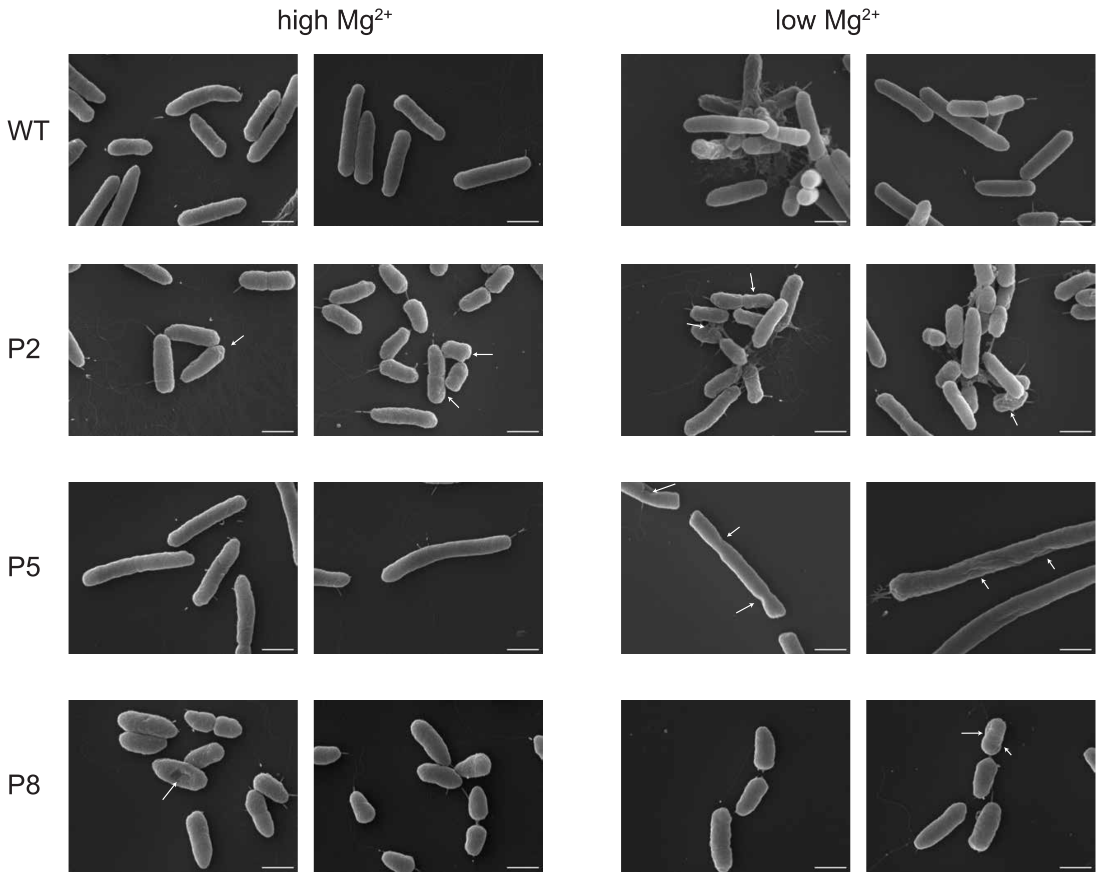


**Fig. S12 Replicates of SEM images of endpoint clones in the high and low $Mg^{2+}$ media.** In the high $Mg^{2+}$ media, P2 and P5, but not P8, have discernible dents or kinks in the cell membrane (white arrows). In low $Mg^{2+}$ media, P5 showed membrane deformation. All three displayed altered cell shapes and lengths compared to WT PAO1. The scale bar indicates 1μm.

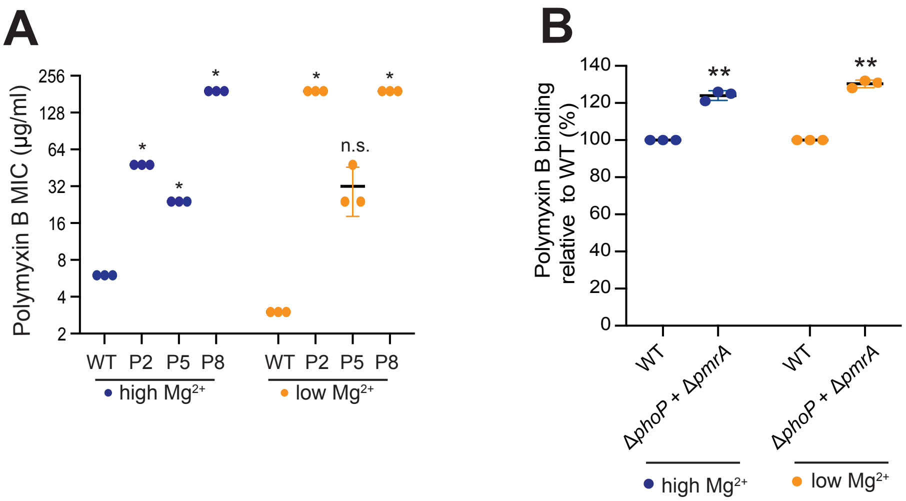


**Fig. S13 Three endpoint clones showed higher polymyxin B MIC than the WT PAO1 (A)** Polymyxin B MIC of endpoint clones in high $Mg^{2+}$ and low $Mg^{2+}$ media. Mean ± std of 3 biological replicates are shown. (* $p<0.05$, Mann-Whitney U test) **(B)** The lack of PhoPQ and PmrAB activity reduced polymyxin B (3µg/mL) binding to the WT PAO1 cells. Dansyl-polymyxin B was used to measure polymyxin B binding to bacteria. Mean ± std of 3 biological replicates are shown. (** $p<0.01$, Dunnett's one-way ANOVA test)

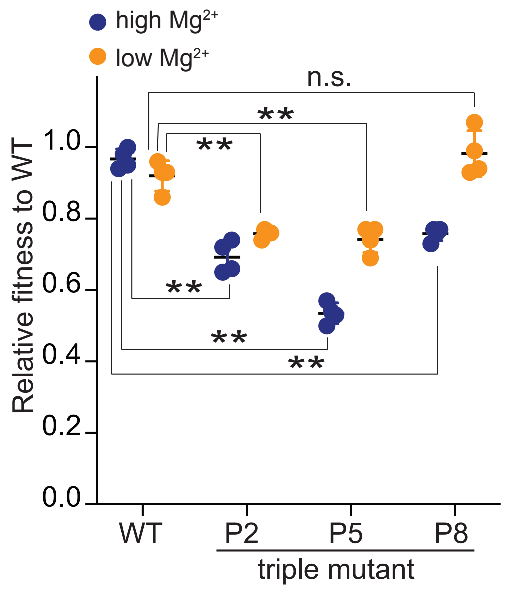


**Fig. S14 Fitness cost of triple mutants in high and low $Mg^{2+}$ conditions**
A competitive fitness assay was used to assess the fitness of three triple mutants relative to WT in high $Mg^{2+}$ (blue) and low $Mg^{2+}$ conditions (orange). Triple mutants of P2 and P5 had reduced fitness in both conditions, while the P8 triple mutant only showed fitness cost in the high $Mg^{2+}$ media. Mean ± std of 4 biological replicates is shown. (**$p < 0.01$, Dunnett's one-way ANOVA test)

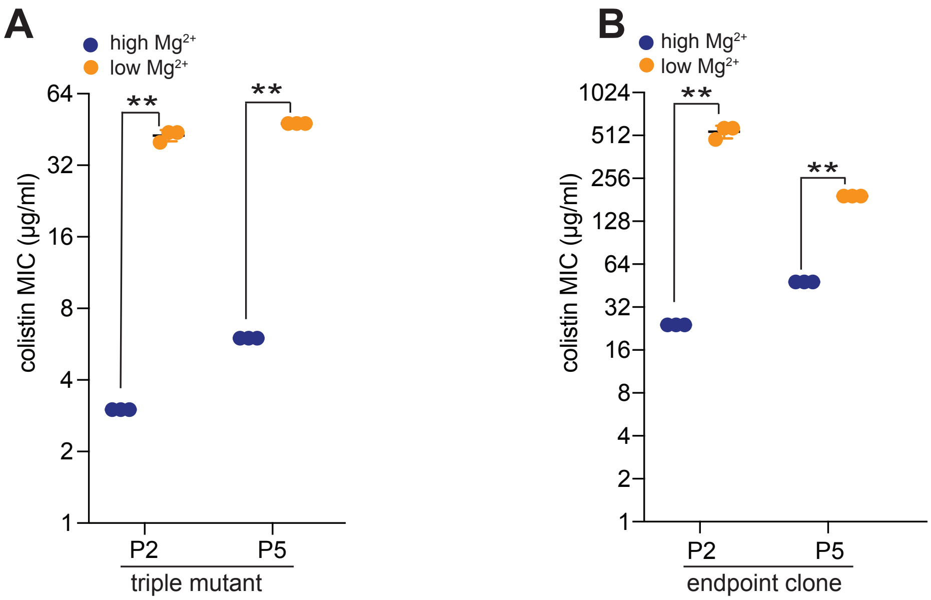


**Fig. S15 P2 and P5 show lower colistin resistance in high $Mg^{2+}$ media**
Colistin resistance of triple mutants (A) and endpoint clones (B) of P2 and P5 was measured by colistin MIC assay in high and low $Mg^{2+}$ media. In both lineages, cells showed significantly lower lower MIC in the high $Mg^{2+}$ media compared to the low $Mg^{2+}$ media. Mean ± std of 3 biological replicates are shown. (**$p < 0.01$, Mann-Whitney U test)

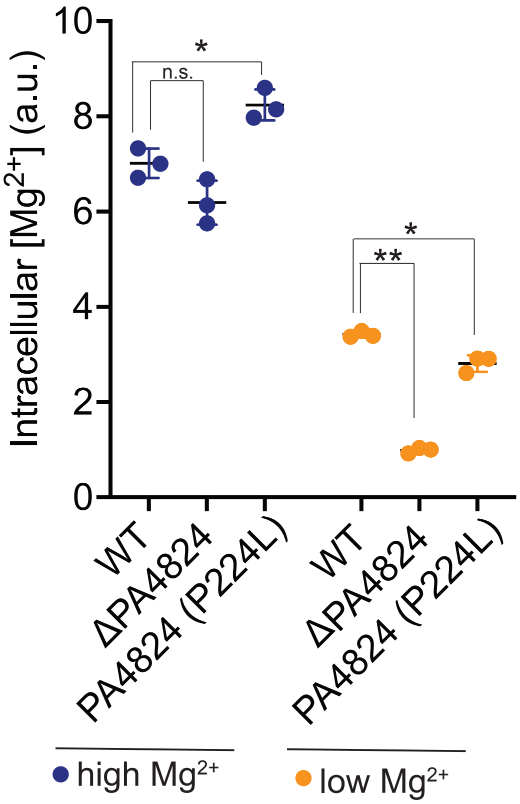


**Fig. S16** ***PA4824*** **mutation has no to mild effects on altering intracellular $Mg^{2+}$ levels**
A $Mg^{2+}$ genetic reporter assay published in the prior study was used to measure the relative intracellular $Mg^{2+}$ levels of WT, Δ*PA4824*, and *PA4824* (P224L) single mutant in high $Mg^{2+}$ and low $Mg^{2+}$ conditions. *PA4824* mutation reconstructed strain had subtle effects on $Mg^{2+}$ levels, indicating this mutation might not impair $Mg^{2+}$ transport function significantly. Mean ± std of 3 biological replicates is shown. (**$p < 0.01$, *$p < 0.05$, Dunnett's one-way ANOVA test)

#### Supplementary Tables

**Table S1. Peak table of all major and minor lipid A species present in MS analysis** Every m/z peak representative of lipid A structure found in FLAT followed by MALDI-TOF MS. C3' and C2' refer to the third and second carbons on the glucosamine on the left side, while C3 and C2 refer to the third and second carbons of the right glucosamine, respectively. Phosphorylation status and aminoarabinose status are not site-specific.

| Calculated m/z | C-3' | C-2' | C-3 | C-2 | Phosphorylation | Aminoarabinose |
|---|---|---|---|---|---|---|
| 1247.6953 | C10(3-OH) | C12(3-OH) | H | C12(3-OH):C12 | Diphosphate | None |
| 1263.693 | C10(3-OH) | C12(3-OH) | H | C12(3-OH):C12(2-OH) | Diphosphate | None |
| 1365.8772 | C10(3-OH) | C12(3-OH):C12 | H | C12(3-OH):C12(2-OH) | Monophosphate | None |
| 1378.7535 | C10(3-OH) | C12(3-OH) | H | C12(3-OH):C12 | Diphosphate | Single |
| 1417.826 | C10(3-OH) | C12(3-OH) | C10(3-OH) | C12(3-OH):C12 | Diphosphate | None |
| 1429.8623 | C10(3-OH) | C12(3-OH):C12 | H | C12(3-OH):C12 | Diphosphate | None |
| 1433.823 | C10(3-OH) | C12(3-OH) | C10(3-OH) | C12(3-OH):C12(2-OH) | Diphosphate | None |
| 1445.8572 | C10(3-OH) | C12(3-OH):C12 | H | C12(3-OH):C12(2-OH) | Diphosphate | None |
| 1461.8522 | C10(3-OH) | C12(3-OH):C12 | H | C12(3-OH):C12(2-OH) | Diphosphate | None |
| 1480.9292 | C10(3-OH) | C12(3-OH):C12 | H | C12(3-OH):C12 | Monophosphate | Single |
| 1485.9249 | C10(3-OH):C16 | C12(3-OH) | H | C12(3-OH):C12 | Diphosphate | None |
| 1496.9241 | C10(3-OH) | C12(3-OH):C12 | H | C12(3-OH):C12(2-OH) | Monophosphate | Single |
| 1501.9199 | C10(3-OH):C16 | C12(3-OH) | H | C12(3-OH):C12(2-OH) | Diphosphate | None |
| 1509.8188 | C10(3-OH) | C12(3-OH) | H | C12(3-OH):C12 | Diphosphate | Double |
| 1548.8842 | C10(3-OH) | C12(3-OH) | C10(3-OH) | C12(3-OH):C12 | Diphosphate | Single |
| 1560.9206 | C10(3-OH) | C12(3-OH):C12 | H | C12(3-OH):C12 | Diphosphate | Single |
| 1564.8812 | C10(3-OH) | C12(3-OH) | C10(3-OH) | C12(3-OH):C12(2-OH) | Diphosphate | Single |
| 1576.9155 | C10(3-OH) | C12(3-OH):C12 | H | C12(3-OH):C12(2-OH) | Diphosphate | Single |
| 1592.9104 | C10(3-OH) | C12(3-OH):C12(2-OH) | H | C12(3-OH):C12(2-OH) | Diphosphate | Single |
| 1599.9954 | C10(3-OH) | C12(3-OH):C12 | C10(3-OH) | C12(3-OH):C12 | Diphosphate | None |
| 1615.9903 | C10(3-OH) | C12(3-OH):C12 | C10(3-OH) | C12(3-OH):C12(2-OH) | Diphosphate | None |
| 1616.9832 | C10(3-OH):C16 | C12(3-OH) | H | C12(3-OH):C12 | Diphosphate | Single |
| 1631.9853 | C10(3-OH) | C12(3-OH):C12(2-OH | C10(3-OH) | C12(3-OH):C12(2-OH) | Diphosphate | None |
| 1632.9781 | C10(3-OH):C16 | C12(3-OH) | H | C12(3-OH):C12(2-OH) | Diphosphate | Single |
| 1668.092 | C10(3-OH):C16 | C12(3-OH):C12 | H | C12(3-OH):C12 | Diphosphate | None |
| 1679.9424 | C10(3-OH) | C12(3-OH) | C10(3-OH) | C12(3-OH):C12 | Diphosphate | Double |
| 1684.0869 | C10(3-OH):C16 | C12(3-OH):C12 | H | C12(3-OH):C12(2-OH) | Diphosphate | None |
| 1691.9561 | C10(3-OH) | C12(3-OH):C12 | H | C12(3-OH):C12 | Diphosphate | Double |

| | | | | | | |
|---|---|---|---|---|---|---|
| 1700.0818 | C10(3-OH):C16 | C12(3-OH):C12(2-OH) | H | C12(3-OH):C12(2-OH) | Diphosphate | None |
| 1707.951 | C10(3-OH) | C12(3-OH):C12 | H | C12(3-OH):C12(2-OH) | Diphosphate | Double |
| 1723.9687 | C10(3-OH) | C12(3-OH):C12(2-OH) | H | C12(3-OH):C12(2-OH) | Diphosphate | Double |
| 1748.0414 | C10(3-OH):C16 | C12(3-OH) | H | C12(3-OH):C12 | Diphosphate | Double |
| 1764.0363 | C10(3-OH):C16 | C12(3-OH) | H | C12(3-OH):C12(2-OH) | Diphosphate | Double |
| 1799.1502 | C10(3-OH):C16 | C12(3-OH):C12 | H | C12(3-OH):C12 | Diphosphate | Single |
| 1815.1452 | C10(3-OH):C16 | C12(3-OH):C12 | H | C12(3-OH):C12(2-OH) | Diphosphate | Single |
| 1831.1401 | C10(3-OH):C16 | C12(3-OH):C12(2-OH) | H | C12(3-OH):C12(2-OH) | Diphosphate | Single |
| 1930.2085 | C10(3-OH):C16 | C12(3-OH):C12 | H | C12(3-OH):C12 | Diphosphate | Double |
| 1946.1849 | C10(3-OH):C16 | C12(3-OH):C12 | H | C12(3-OH):C12(2-OH) | Diphosphate | Double |
| 1962.1983 | C10(3-OH):C16 | C12(3-OH):C12(2-OH) | H | C12(3-OH):C12(2-OH) | Diphosphate | Double |

### Table S2. Strains used in this study

| strain name | specie name | genotype | lab source | Additional Information |
|---|---|---|---|---|
| YPH-b1 | *P. aeruginosa* PAO1 | WT | PAO1 master strain from the Dandekar lab, UW | |
| YPH-b3 | *P. aeruginosa* PAO1 | WT mCherry (at neutral site) | Generated in this study | |
| YPH-b304 | *P. aeruginosa* PAO1 | PA4824(P224L) in PAO1 | Generated in this study | Mutation reconstruction from P2 |
| YPH-b370 | *P. aeruginosa* PAO1 | htrB2(R294H) in PAO1 | Generated in this study | Mutation reconstruction from P2 |
| YPH-b371 | *P. aeruginosa* PAO1 | pOprH(A to G) in PAO1 | Generated in this study | Mutation reconstruction from P2 |
| YPH-b385 | *P. aeruginosa* PAO1 | PA4824(P224L) + pOprH(A to G) in PAO1 | Generated in this study | Mutation reconstruction from P2 |
| YPH-b389 | *P. aeruginosa* PAO1 | PA4824(P224L) + htrB2(R294H) + pOprH(A to G) triple mutation in PAO1 | Generated in this study | Mutation reconstruction from P2 |
| YPH-b365 | *P. aeruginosa* PAO1 | htrB2 (R170W) in PAO1 | Generated in this study | Mutation reconstruction from P5 |
| YPH-b387 | *P. aeruginosa* PAO1 | lpxO2 (D163A) in PAO1 | Generated in this study | Mutation reconstruction from P5 |
| YPH-b368 | *P. aeruginosa* PAO1 | PA4824(P224L) + htrB2(R170W) in PAO1 | Generated in this study | Mutation reconstruction from P5 |
| YPH-b418 | *P. aeruginosa* PAO1 | PA4824 (P224L) + lpxO2(D163A) + htrB2(R170W) triple mutation in PAO1 | Generated in this study | Mutation reconstruction from P5 |
| YPH-b421 | *P. aeruginosa* PAO1 | pOprH (A to C) in PAO1 | Generated in this study | Mutation reconstruction from P8 |
| YPH-b419 | *P. aeruginosa* PAO1 | OprH(A to C) + lpxA (G170C) in PAO1 | Generated in this study | Mutation reconstruction from P8 |
| YPH-b420 | *P. aeruginosa* PAO1 | PA4824 (P224L) + OprH (A to C) + lpxA (G170C) triple mutation in PAO1 | Generated in this study | Mutation reconstruction from P8 |

| | | | | |
|---|---|---|---|---|
| YPH-b429 | *P. aeruginosa* PAO1 | WT htrB2 in P2 | Generated in this study | |
| YPH-b451 | *P. aeruginosa* PAO1 | WT pOprH in P2 | Generated in this study | |
| YPH-b435 | *P. aeruginosa* PAO1 | WT PA4824 in P2 | Generated in this study | |
| YPH-b423 | *P. aeruginosa* PAO1 | WT PA4824 in P5 | Generated in this study | |
| YPH-b424 | *P. aeruginosa* PAO1 | WT htrB2 in P5 | Generated in this study | |
| YPH-b450 | *P. aeruginosa* PAO1 | WT lpxO2 in P5 | Generated in this study | |
| YPH-b426 | *P. aeruginosa* PAO1 | WT PA4824 in P8 | Generated in this study | |
| YPH-b425 | *P. aeruginosa* PAO1 | WT OprH in P8 | Generated in this study | |
| YPH-b470 | *P. aeruginosa* PAO1 | OprH deletion in P2 | Generated in this study | |
| YPH-b458 | *P. aeruginosa* PAO1 | OprH deletion in P8 | Generated in this study | |
| YPH-b468 | *P. aeruginosa* PAO1 | phop deletion in P2 | Generated in this study | |
| YPH-b462 | *P. aeruginosa* PAO1 | phop deletion in P8 | Generated in this study | |
| YPH-b481 | *P. aeruginosa* PAO1 | phoP deletion + pmrA deletion in P2 | Generated in this study | |
| YPH-b479 | *P. aeruginosa* PAO1 | phoP deletion + pmrA deletion in P5 | Generated in this study | |
| YPH-b472 | *P. aeruginosa* PAO1 | phoP deletion + pmrA deletion in P8 | Generated in this study | |
| YPH-b483 | *P. aeruginosa* PAO1 | PA4824(P224L) + htrB2(R294H) + pOprH(A to G) triple mutation in PAO1 + phoP deletion | Generated in this study | Mutation reconstruction from P2 |
| YPH-b476 | *P. aeruginosa* PAO1 | PA4824 (P224L) + lpxO2(D163A) + htrB2(R170W) triple mutation in PAO1 + phoP deletion | Generated in this study | Mutation reconstruction from P5 |
| YPH-b477 | *P. aeruginosa* PAO1 | PA4824 (P224L) + OprH (A to C) + lpxA (G170C) triple mutation in PAO1 + phoP deletion | Generated in this study | Mutation reconstruction from P8 |
| YPH-b485 | *P. aeruginosa* PAO1 | PA4824(P224L) + htrB2(R294H) + pOprH(A to G) triple mutation in PAO1 + phoP deletion + pmrA deletion | Generated in this study | Mutation reconstruction from P2 |
| YPH-b487 | *P. aeruginosa* PAO1 | PA4824 (P224L) + lpxO2(D163A) + htrB2(R170W) triple mutation in PAO1 + phoP deletion + pmrA deletion | Generated in this study | Mutation reconstruction from P5 |
| YPH-b486 | *P. aeruginosa* PAO1 | PA4824 (P224L) + OprH (A to C) + lpxA (G170C) triple mutation in PAO1 + phoP deletion + pmrA deletion | Generated in this study | Mutation reconstruction from P8 |
| YPH-b390 | *P. aeruginosa* PAO1 | phop deletion in P5 | Generated in this study | |
| YPH-b465 | *P. aeruginosa* PAO1 | phop deletion in WT PAO1 | Generated in this study | |
| YPH-b471 | *P. aeruginosa* PAO1 | pmrA deletion in WT PAO1 | Generated in this study | |
| YPH-b474 | *P. aeruginosa* PAO1 | phoP deletion + pmrA deletion in WT PAO1 | Generated in this study | |
| YPH-b504 | *P. aeruginosa* PAO1 | pmrA deletion in P2 | | |
| YPH-b500 | *P. aeruginosa* PAO1 | pmrA deletion in P5 | | |
| YPH-b499 | *P. aeruginosa* PAO1 | pmrA deletion in P8 | | |
| YPH-y1 | *C. albicans* | WT prototroph (SC5314) | From Dr. Alistair Brown | |

#### Table S3. Primers used in this study

| Primer number | Primer name | Primer sequence | Note |
|---|---|---|---|
| YPH_o1 | pEXG2_linear _R_v3 | GCTTGCTTTACATTTATGCTTCC | Gibson Assembly for making pEXG2-based allelic exchange plasmid |
| YPH_o2 | pEXG2_linear _F_v3 | ACTCTAGAGGATCCCCGG | Gibson Assembly for making pEXG2-based allelic exchange plasmid |
| YPH_401 | Q5_PA4824_P 224L_F | GCAGGCCGGActgCGGCTGCACC | for making point mutation using Q5 mutagenesis kit |
| YPH_402 | Q5_PA4824_P 224L_R | CAGCCGCTGTCGCCGGCC | for making point mutation using Q5 mutagenesis kit |
| YPH_403 | PA4824_CDS_ seq_F | GAGCTGAAGACACCGCTG | for sequecning PA4824 P224L mutation |
| YPH_404 | PA4824_3UT R_seq_R | AGGGAACGGGCTCATTGA | for sequecning PA4824 P224L mutation |
| YPH_415 | pEXG2-lpxO2-F | CACACATTATACGAGCCGGAAGCATAAATGTAAAG CAAGCGCAGGCGCGTCCCTTC | Gibson Assembly for making pEXG2-based allelic exchange plasmid |
| YPH_416 | pEXG2-lpxO2-R | GGTACCGAATTCGAGCTCGAGCCCGGGGATCCTCTA GAGTAACTGAAGGCCGAGGAGC | Gibson Assembly for making pEXG2-based allelic exchange plasmid |
| YPH_419 | pEXG2-htrB2-F | CACACATTATACGAGCCGGAAGCATAAATGTAAAG CAAGCTCGCGGGCGCCC | Gibson Assembly for making pEXG2-based allelic exchange plasmid |
| YPH_420 | pEXG2-htrB2-R | GGTACCGAATTCGAGCTCGAGCCCGGGGATCCTCTA GAGTCGGCGCTGGCGCA | Gibson Assembly for making pEXG2-based allelic exchange plasmid |
| YPH_421 | pEXG2-poprH-F | CACACATTATACGAGCCGGAAGCATAAATGTAAAG CAAGCGAAGCGGATCGCGGC | Gibson Assembly for making pEXG2-based allelic exchange plasmid |
| YPH_422 | pEXG2-poprH-R | GGTACCGAATTCGAGCTCGAGCCCGGGGATCCTCTA GAGTGCGGGGTCATTTAGAACTTG | Gibson Assembly for making pEXG2-based allelic exchange plasmid |
| YPH_429 | Q5_htrB2_R17 0W_F | TGGCCGCGAGtggCACAACCTCG | for making point mutation using Q5 mutagenesis kit |
| YPH_430 | Q5_htrB2_R17 0W_R | CGGCGCTGGACGTAGTCGAAC | for making point mutation using Q5 mutagenesis kit |
| YPH_431 | Q5_htrB2_R29 4H_F | GGCGCACCGGcatTTCAAGACCC | for making point mutation using Q5 mutagenesis kit |
| YPH_432 | Q5_htrB2_R29 4H_R | CACAGGTACTGCTCGGGC | for making point mutation using Q5 mutagenesis kit |
| YPH_438 | lpxO2_CDS_F | GCTTCTACCTGACCTGGT | for sequencing lpxO2 mutation |
| YPH_439 | lpxO2_CDS_R | GGCGGAGGATTCGGTAG | for sequencing lpxO2 mutation |
| YPH_442 | htrB2_CDS_F | CATTTCACCACCCTGGAG | for sequencing htrB2 mutation |
| YPH_443 | htrB2_CDS_R | CCCGGAAAGGATCAGGC | for sequencing htrB2 mutation |
| YPH_456 | newQ5_lpxO2 _D163A_F | CCGCACCGCGgcccCTTCGGCGGC | for making point mutation using Q5 mutagenesis kit |
| YPH_457 | newQ5_lpxO2 _D163A_R | ATTCAGGTGGCTGCCGCC | for making point mutation using Q5 mutagenesis kit |
| YPH_460 | Q5_pOprH_G_F | GCAAGCGTTCcggGGCGGTTCAG | for making point mutation using Q5 mutagenesis kit |
| YPH_461 | Q5_pOprH_G_R | TGAACCCGGGCTGAA | for making point mutation using Q5 mutagenesis kit |
| YPH_476 | pOprH-F | CAGTAGCTGGTACATCCAGA | for sequencing poprH mutation |
| YPH_477 | pOprH-R | TCTTGCTCAGCTCCTGCA | for sequencing poprH mutation |
| YPH_478 | pEXG2-phoP-F | ATTCCACACATTATACGAGCCGGAAGCATAAATGTA AAGCAAGCCACCTGGGGCATCCGC | Gibson Assembly for making pEXG2-based deletion plasmid |
| YPH_479 | pEXG2-phoP-R | TAAGGTACCGAATTCGAGCTCGAGCCCGGGGATCCT CTAGAGTCAGGTAGAGCTGCTCGC | Gibson Assembly for making pEXG2-based deletion plasmid |
| YPH_480 | phoP-DOWN_F | CCGGCCGCATCTCATCCGGAGGAACCTCTCCGTTCC CTGCGCATC | Gibson Assembly for making pEXG2-based deletion plasmid |
| YPH_481 | phoP-UP-R | ATCAGACGGATGCGCAGGGAACGGAGAGGTTCCTC CGGATGAGAT | Gibson Assembly for making pEXG2-based deletion plasmid |
| YPH_496 | phoP_check_F | TGCAGGCCGGTATCCT | for checking phoP deletion |
| YPH_497 | phoP_check_R | CTGGTCGTAGATATAGCCGAG | for checking phoP deletion |
| YPH_498 | pOprH_C_F | CCCGGGTTCCGCAAGCGTTCAGGGGCGG | for making point mutation using Q5 mutagenesis kit |

| | | | |
|---|---|---|---|
| YPH_499 | pOprH_C_R | CTTGCGGAACCCGGGCTGAACGACTCG | for making point mutation using Q5 mutagenesis kit |
| YPH_506 | phoP_FWD | TGA AAC TGC TGG TAG TGG AAG | for QPCR |
| YPH_507 | phoP_REV | TAT TCG CTG ACC CGG TAG A | for QPCR |
| YPH_510 | pmrB_FWD | CTC ATC GAC GAA CTC AAC CTC | for QPCR |
| YPH_511 | pmrB_REV | TGG CGT GCG GAT TTC AT | for QPCR |
| YPH_524 | arnc_FWD | AGT ACC TTC ATC CCG ATC CT | for QPCR |
| YPH_525 | arnc_REV | CAT CGC GCT GTA TTT CGA TTC | for QPCR |
| YPH_563 | pagP_F | TTTCGACAGTGACAGCTACC | for QPCR |
| YPH_564 | pagP_R | GGGATCTTGTCGCGGTATT | for QPCR |
| YPH_569 | phoQ_check_F | AGTTCTAAATGACCGCGCA | for QPCR |
| YPH_570 | phoQ_check_R | AGCCAGATCGCAGCGATG | for QPCR |
| YPH_581 | rplU_FWD | CAAAGTCACCGAAGGCGAAT | for QPCR |
| YPH_582 | rplU_REV | ACGCTTCATGTGGTGCTTAC | for QPCR |
| YPH_571 | pmrA_FWD | GACATCCTGCGCAACCT | for QPCR |
| YPH_572 | pmrA_REV | GATCGAAGGGCTTGGTCA | for QPCR |
| YPH_500 | pEXG2-pmrAdel-F | AATTCCACACATTATACGAGCCGGAAGCATAAATGT AAAGCAAGCGCAACCCCGAGCGCT | Gibson Assembly for making pEXG2-based deletion plasmid |
| YPH_501 | pEXG2-pmrAdel-R | TTAAGGTACCGAATTCGAGCTCGAGCCCGGGGATCC TCTAGAGTCATGGTGTGCTGGCGG | Gibson Assembly for making pEXG2-based deletion plasmid |
| YPH_502 | pmrA-DOWN-F | ACTGAAACGAGGCTGCCCCCGCCGCACCACC | Gibson Assembly for making pEXG2-based deletion plasmid |
| YPH_503 | pmrA-UP-R | GGTGGTGCGGCGGGGGCAGCCTCGTTTCAGT | Gibson Assembly for making pEXG2-based deletion plasmid |
